# UC-associated autoantibodies to αvβ6 inhibit mucosal TGFβ activation and predispose to intestinal inflammation

**DOI:** 10.64898/2026.03.12.709585

**Authors:** Kayla J. Fasano, Anna E. Yoshida, Jane F. Madden, Kelsey E. Mauk, Lin Wei Tung, Thomas H. Edwards, Donna M. Shows, Caroline Stefani, David G. Kugler, Sheila Scheiding, Adarsh B. Manjunath, Megan E. Smithmyer, Oliver J. Harrison, Cate Speake, James D. Lord, Adam Lacy-Hulbert

## Abstract

Ulcerative colitis (UC) is characterized by epithelial barrier dysfunction and dysregulated mucosal immune responses; however, the mechanisms driving disease onset remain poorly defined. Autoantibodies against the epithelial-restricted integrin αvβ6 are a highly specific biomarker of UC that can precede clinical diagnosis by up to 10 years. Because αvβ6 activates TGFβ at epithelial surfaces, we hypothesized that UC-associated αvβ6 autoantibodies inhibit mucosal TGFβ activation and disrupt epithelial homeostasis. We showed that αvβ6 autoantibodies were enriched in UC and that IgG from autoantibody-positive individuals inhibited αvβ6–dependent activation of TGFβ. αvβ6 blockade dampened TGFβ signaling and altered differentiation–associated gene programs in human intestinal epithelial cells. In mice, deletion of αv caused expansion of inflammation-associated goblet cells in the colon and changes in intestinal immune cells. Using a novel mouse model, we showed that αvβ6-specific autoantibody disrupted epithelial–immune crosstalk and increased susceptibility to DSS colitis. Together, these findings establish anti-αvβ6 autoantibodies as active inhibitors of epithelial TGFβ signaling, constituting a *de facto* anti-cytokine response, rather than passive biomarkers. By linking preclinical seropositivity to impaired epithelial signaling and heightened susceptibility to colitis, this work identifies epithelial αvβ6–dependent TGFβ activation as a pathway that may be leveraged to modify disease risk or limit disease severity.

**One Sentence Summary:** UC-associated autoantibodies impair epithelial TGFβ activation, alter mucosal homeostasis, and predispose to colitis.

## Introduction

Ulcerative colitis (UC) is a chronic inflammatory bowel disease (IBD) of the colorectum characterized by relapsing inflammation of the mucosal layer and progressive disruption of gut barrier function. The incidence and prevalence of UC continue to rise worldwide, posing a substantial and growing global health burden (*1*). Clinically, UC is characterized by local and systemic manifestations that significantly impair quality of life. Symptoms include abdominal pain, diarrhea, and rectal bleeding, and many of those diagnosed with UC will require long-term immunosuppressive therapy or colectomy. Although current therapies can induce remission in a subset of individuals, a substantial proportion fail to respond or lose response over time (*2, 3*), underscoring the need to better understand early pathogenic mechanisms of disease and identify new therapeutic targets.

A central feature of UC is breakdown of the intestinal mucosal barrier which normally serves as a tightly regulated interface between luminal microbiota and the host immune system. The mucosal surface of the colon is comprised primarily of intestinal epithelial cells (IECs), including absorptive enterocytes, secretory and sensory cells, and specialized resident immune cells, which together form a physical and immunological barrier (*4*). In UC, disruption of epithelial integrity leads to increased permeability, aberrant immune activation and sustained inflammation, contributing to exacerbation of disease (*4–6*).

Increasing evidence suggests that UC, like other autoimmune diseases, is preceded by a preclinical phase characterized by subclinical immune activation and measurable serologic changes years before diagnosis (*7*). In diseases such as rheumatoid arthritis and systemic lupus erythematosus, disease-specific autoantibodies arise long before clinical onset and are now understood to participate directly in disease pathogenesis, rather than serving solely as biomarkers (*8*). However, UC has not traditionally been viewed as an antibody-mediated autoimmune disease, and the functional contribution of preclinical autoantibodies remains poorly defined. Recent reports have identified autoantibodies against the epithelial cell-specific integrin αvβ6 as biomarkers for UC that can precede clinical diagnosis by up to 10 years (*9–15*). While these findings establish αvβ6 autoantibodies as highly specific biomarkers of preclinical disease, whether they actively impact epithelial function or simply reflect secondary immune activation remains unknown. Notably, αvβ6 has been shown to play a role in the maintenance of barrier integrity, suggesting its function may be mechanistically important in the development of UC (*16, 17*).

αvβ6 belongs to the αv family of integrins which form cell surface heterodimers composed of an α and a β subunit (*18*). αvβ6 is expressed exclusively by epithelial cells and binds to an Arginine-Glycine-Aspartic acid (RGD) amino acid motif present on a variety of extracellular matrix proteins. One of its major functions is the ability to activate the cytokine TGFβ (*16, 17*). TGFβ is a multifunctional cytokine produced as 3 isoforms: TGF-β1, TGF-β2 and TGF-β3. Of these, TGF-β1 (henceforward referred to as TGFβ) is the most abundantly expressed in the intestine where it is produced by and signals to multiple cell types, including epithelial and immune cells. TGFβ is made as an inactive precursor, where the active cytokine is shielded by latency-associated peptide (LAP). αvβ6 activates TGFβ by binding to the RGD sequence contained within LAP, initiating a mechanical force that pulls the latency complex apart, exposing TGFβ’s receptor-binding site (*16, 17*). Importantly, αvβ6-mediated activation does not appear to fully release TGFβ from the latency complex, requiring close cell-to-cell contact for signaling (*19, 20*).

A large body of work has shown a critical role of the TGFβ signaling pathway in the pathogenesis of IBD. Studies in individuals with IBD have shown decreased phosphorylation of downstream targets of TGFβ, SMAD3/SMAD4 and increased expression of the negative regulator of TGFβ, SMAD7, in mucosal samples (*21*). Furthermore, genetic variants in the SMAD family of genes are associated with the development of UC (*22*). In experimental mouse models, knockout of TGFβ signaling components globally or in specific immune cell subsets induces spontaneous colitis or increases susceptibility to DSS colitis (*23–26*). Highlighting the importance of intrinsic TGFβ signaling in the intestinal epithelium, loss of TGFβ receptor (*Tgfbr2*) function specifically in IECs similarly increases susceptibility to DSS colitis and impairs intestinal wound healing (*27–29*). Given that impaired TGFβ signaling is strongly linked to intestinal inflammation, understanding how latent TGFβ is locally activated within the colonic epithelium and how this process may be perturbed in UC is critical.

Although αvβ6 autoantibodies have emerged as specific biomarkers of UC, fundamental questions remain regarding their functional properties and relevance to disease pathophysiology. Recently, rare loss-of-function variants in the genes encoding αv (*ITGAV*) and β6 (*ITGB6*) have been associated with severe early onset colitis and intestinal inflammation (*30, 31*). Given these findings, the central role of αvβ6 in activation of TGFβ, and the importance of epithelial TGFβ signaling in barrier maintenance and repair, we hypothesized that UC-associated αvβ6 autoantibodies inhibit αvβ6-mediated TGFβ activation, disrupt epithelial homeostasis, and establish a pre-colitis state that precedes and predisposes to the development of disease. To test this, we integrated analyses of samples from individuals with UC and functional assays of αvβ6-dependent TGFβ activation, transcriptional profiling in human intestinal epithelial cells, and complementary *in vivo* models of loss of epithelial αvβ6 function. Together, our findings define a mechanistic link between anti-αvβ6 autoantibodies, impaired mucosal TGFβ activation, and increased susceptibility to colitis.

## Results

### Anti-αvβ6 IgG and IgA are detected in UC

To confirm recent reports that anti-αvβ6 autoantibodies are strongly associated with UC, we analyzed serum and plasma from individuals with UC (n=194) and healthy subjects (HS) with no family history of autoimmune disease (n=338). (**Table 1**). We detected both IgG and IgA binding to αvβ6 in UC samples by ELISA, but not in the majority of HS (**Fig 1A,B**). In UC, levels of IgG and IgA autoantibodies were significantly correlated (r=0.61, p<0.001; **Supp Fig 1A**). Total IgG and IgA were significantly but modestly elevated in UC compared to HS (**Supp Fig 1B**).

**Fig. 1.**
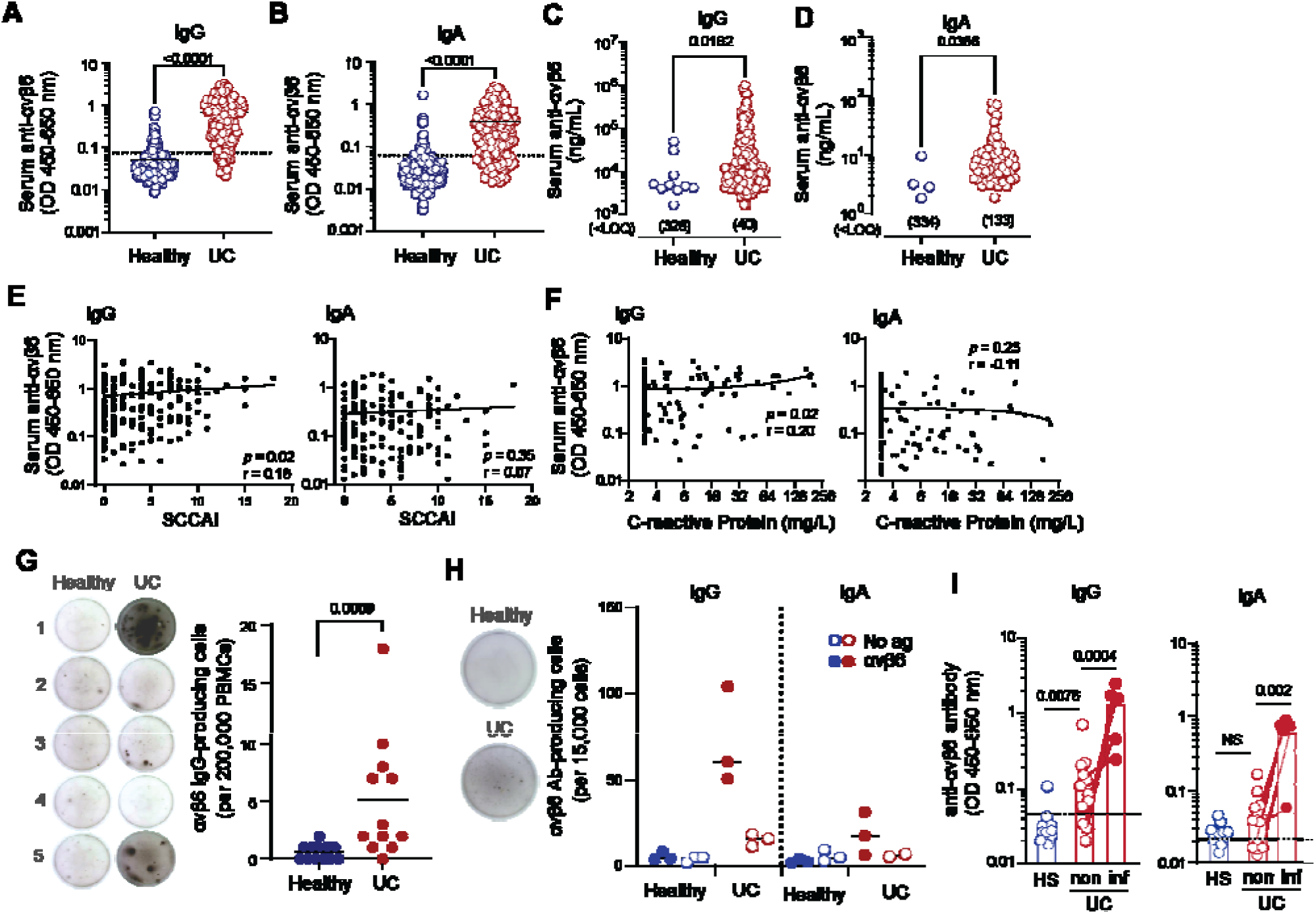
Anti-αvβ6 antibody levels, clinical parameters, and αvβ6-specific B cells in UC and healthy subjects. (**A-B**) Serum anti-αvβ6 IgG (**A**) and IgA (**B**) quantified by ELISA in healthy subjects (HS; n = 338) and ulcerative colitis (UC; n = 194). Dashed lines indicate OD thresholds for autoantibody positivity, determined by receiver operating characteristic (ROC) curve analysis using the highest Youden’s index. (**C-D**) Concentrations of anti-αvβ6 IgG (HS, n=10; UC n=153) (**C**) and IgA (HS, n=4; UC n=61) (**D**) in sera were interpolated from monoclonal antibody standard curves; samples below the lower limit of quantification (LLOQ) are indicated and were excluded from concentration-based statistical analysis. (**E-F**) Correlation between serum anti-αvβ6 IgG (left) or IgA (right) OD values and Simple Clinical Colitis Activity Index (SCCAI) scores (n=171) (**E**) or serum C-reactive protein (CRP) levels (n=132) (**F**) in UC donors. (**G**) Representative IgG ELISPOT images (left) and quantification (right) of αvβ6-specific IgG-producing cells in peripheral blood mononuclear cells. (**H**) Representative IgG ELISPOT images (left) and quantification (right) of αvβ6-specific IgG and IgA-producing cells in colonic biopsies. Wells coated without αvβ6 served as negative controls. (**I, J**) Anti-αvβ6 IgG (**I**) and IgA (**J**) levels measured in supernatants from overnight-cultured colonic biopsies; dotted lines indicate lower limits of detection (LLOD). Data are shown as individual data points with medians (**A-D, G-J**) or with best-fit lines for visualization purposes only (**E-F**). Two-group comparisons were performed using two-tailed Mann-Whitney tests (**A-D, G, I, J**). Correlations were assessed using two-tailed Spearman rank correlation (**E-F**).

**Table 1.** Demographic and clinical characteristics of study cohorts. Baseline demographic and clinical features of participants included in the ulcerative colitis (UC) and healthy subject (HS) cohorts. Values are presented as mean ± standard deviation (SD) for continuous variables and percentage of participants for categorical variables. Body mass index (BMI), Simple Clinical Colitis Activity Index (SCCAI), C-reactive protein (CRP). Disease-specific variables (duration, SCCAI, CRP, and medication use at the time of blood draw) were collected for the UC cohort only and are therefore listed as not applicable (n/a) for the HS cohort. Medication categories are not mutually exclusive; percentages reflect the proportion of UC participants receiving each therapy at the time of sample collection.

| Characteristic | UC Cohort<br>(n=194) | HS Cohort<br>(n=338) |
| --- | --- | --- |
| **Mean age ±SD** | 42.8 ±15 | 51.2 ±16.8 |
| **Female (%)** | 47.9% | 73.2% |
| **Race** |  |  |
| Native American | 2.1% | 0.3% |
| Asian | 4.8% | 5.3% |
| Black | 1.6% | 2.4% |
| White | 83.5% | 86.7% |
| Multiple | 2.7% | 4.4% |
| Not reported | 5.3% | 1.5% |
| **Hispanic or Latino** |  |  |
| Yes | 6.9% | 4.4% |
| No | 86.7% | 93.2% |
| Not reported | 6.4% | 2.4% |
| **Mean weight in kg ±SD** | 78.4 ±21.0 | 74.0 ±18.3 |
| **Mean BMI ±SD** | 26.3 ±5.7 | 26.0 ±5.5 |
| **UC duration in years ±SD** | 9.5 ±10.1 | n/a |
| **SCCAI ±SD** | 4.4 ±3.9 | n/a |
| **Mean C reactive protein ±SD** | 16.3 ±35.6 | n/a |
| **Medication use at draw** |  |  |
| Glucocorticoids | 24.2% | n/a |
| Aminosalicylates | 52.2% | n/a |
| Thiopurines | 12.6% | n/a |
| Anti-TNF | 18.7% | n/a |
| Vedolizumab | 6.6% | n/a |
| Ustekinumab | 0.5% | n/a |
| Janus kinase inhibitor | 4.4% | n/a |
| None | 23.6% | n/a |

Anti-αvβ6 autoantibody levels strongly discriminated those with UC from HS (area under the receiver operator curve [AUC] of 0.959 for IgG and 0.899 for IgA; **Supp Fig 1C**), and using thresholds derived from this analysis, 173 (89.2%) of UC donors were positive for anti-αvβ6 IgG, and 148 (76.2%) for IgA, compared with only 39 (11.5%) and 35 (10.4%) of HS. To estimate relative autoantibody abundance, we also calculated equivalent αvβ6 binding units compared with monoclonal human anti-αvβ6 IgG or IgA. By this metric, very few (≤10) HS samples exhibited detectable anti-αvβ6, and at a significantly lower concentration than UC samples (**Fig 1 C,D**).

Antibodies to the related integrin αvβ3 have been reported in some UC donors (*9*) and isolated anti-αvβ6 autoantibodies can show cross reactivity with αvβ3 (*14*). To determine whether anti-αvβ6 seropositivity reflected broader anti-αv integrin reactivity, we screened a subset of samples for anti-αvβ3 antibodies. Anti-αvβ3 OD values were significantly higher in UC compared with HS (**Supp Fig 1D**); however, substantial overlap between groups was observed. Importantly, in UC, anti-αvβ3 IgG levels did not correlate with anti-αvβ6 IgG levels (r=0.11, *p*<0.35; **Supp Fig 1E**), while a modest correlation between αvβ6 and αvβ3 IgG was observed in healthy subjects (r=0.34, *p*<0.0001; **Supp Fig 1F**). Together, these findings indicate that UC αvβ6 autoantibodies reflect a selective immune response against αvβ6 integrin rather than a manifestation of broad anti-αv autoreactivity.

Prior studies examining whether anti-αvβ6 autoantibody levels correlate with disease severity have yielded mixed results (*9, 32, 33*). In our cohort, autoantibody levels demonstrated weak or no correlation with clinical disease activity (SCCAI; **Fig 1E**) or systemic inflammation (C-reactive protein; **Fig 1F**) and did not meaningfully associate with fecal calprotectin levels (**Suppl Fig 1G**). Among those with UC, duration of disease, age, and body mass index did not correlate with autoantibody levels, and there was no significant difference by sex (**Supp Fig 1H-K**).

Furthermore, the majority of individuals that had undergone colectomy remained seropositive (**Supp Fig 1L)**. As has been previously reported (*34–36*), IgG autoantibodies were significantly higher in a small subset of people with UC and concomitant primary sclerosing cholangitis (PSC), compared with non-PSC UC (**Suppl Fig 1M)**. Collectively, these data indicate that anti-αvβ6 autoantibody levels do not consistently track with inflammatory activity or most clinical severity metrics in established UC.

### **α**v**β**6-specific B cells and autoantibodies are detected in UC tissue

Previous studies have focused on autoantibodies present in blood circulation, but it remains unclear where autoreactive B cells reside and whether autoantibodies are produced at sites of inflammation in the colon. To answer this question, we developed an ELISPOT assay to detect αvβ6-specific B cells. We first tested for the presence of αvβ6-specific memory B cells in circulation using PBMCs from clinically confirmed UC and age-matched HS pre-activated with R848 and IL-2 (*37*). αvβ6-specific B cells were detected exclusively in UC PBMCs (**Fig 1G**).

Given our ability to detect αvβ6-specific B cells in peripheral blood, we next assayed lamina propria (LP) cells from colon biopsies of people with and without UC. To focus on antibody-secreting plasma cells, LP cells were not activated prior to the ELISPOT assay. We detected anti-αvβ6 IgG-producing B cells exclusively in UC (**Fig 1H**). Anti-αvβ6 IgA-producing cells were also detected in the LP, but at lower numbers than IgG-producing cells. To further confirm that autoantibody-producing cells are present in UC LP, we measured autoantibodies in supernatants from overnight cultures of colon biopsies. Both IgG and IgA anti-αvβ6 autoantibodies were produced by biopsies from UC but not HS. Furthermore, in a subset of samples with paired biopsies from the same UC donor, both IgG and IgA autoantibodies were elevated in inflamed (defined by gross endoscopic inflammation at time of biopsy) relative to uninflamed regions of the colon (**Fig 1I**). Together, these findings indicate that αvβ6-specific B cells are present both in circulation and in colonic tissue and suggest that local autoantibody production is increased during inflammation.

### IgG from autoantibody positive UC inhibits activation of TGF**β**

αvβ6 binds to a range of extracellular matrix-associated proteins through a conserved Arginine-Glycine-Aspartic acid (RGD) amino acid motif. Prior work has shown that UC autoantibodies block the ability of αvβ6 to bind to the RGD-containing ligand fibronectin (*9, 14*). However, a major physiological role for αvβ6 is to activate TGFβ by binding to a another RGD-containing ligand, LAP, on TGFβ (*16, 17*). We therefore sought to determine whether UC autoantibodies inhibited this function of αvβ6. We first tested whether serum from 29 UC donors who exhibited the highest IgG autoantibody titers could inhibit αvβ6 binding to latent TGFβ (L-TGFβ) using HT-29 colonic epithelial cells that endogenously express αvβ6 (*9*). HT-29 cells adhered to plates coated with recombinant L-TGFβ, and this was inhibited by pretreatment with an αvβ6-specific monoclonal antibody, 3G9, that blocks ligand binding (*38, 39*). Serum from 10 of 29 UC inhibited cell adhesion to L-TGFβ, whereas serum from HS with no detectable anti-αvβ6 antibodies had no effect (**Fig 2A**). Of note, sera that inhibited adhesion to L-TGFβ also inhibited binding to fibronectin, consistent with antibodies inhibiting integrin binding to the canonical RGD ligand (**Supp Fig 2A,B**). Anti-αvβ6 IgG and IgA titers did not relate to inhibitory function, suggesting function-blocking antibodies may comprise a variable fraction of the polyclonal anti-αvβ6 repertoire (**Fig 2B).**

**Fig. 2.**
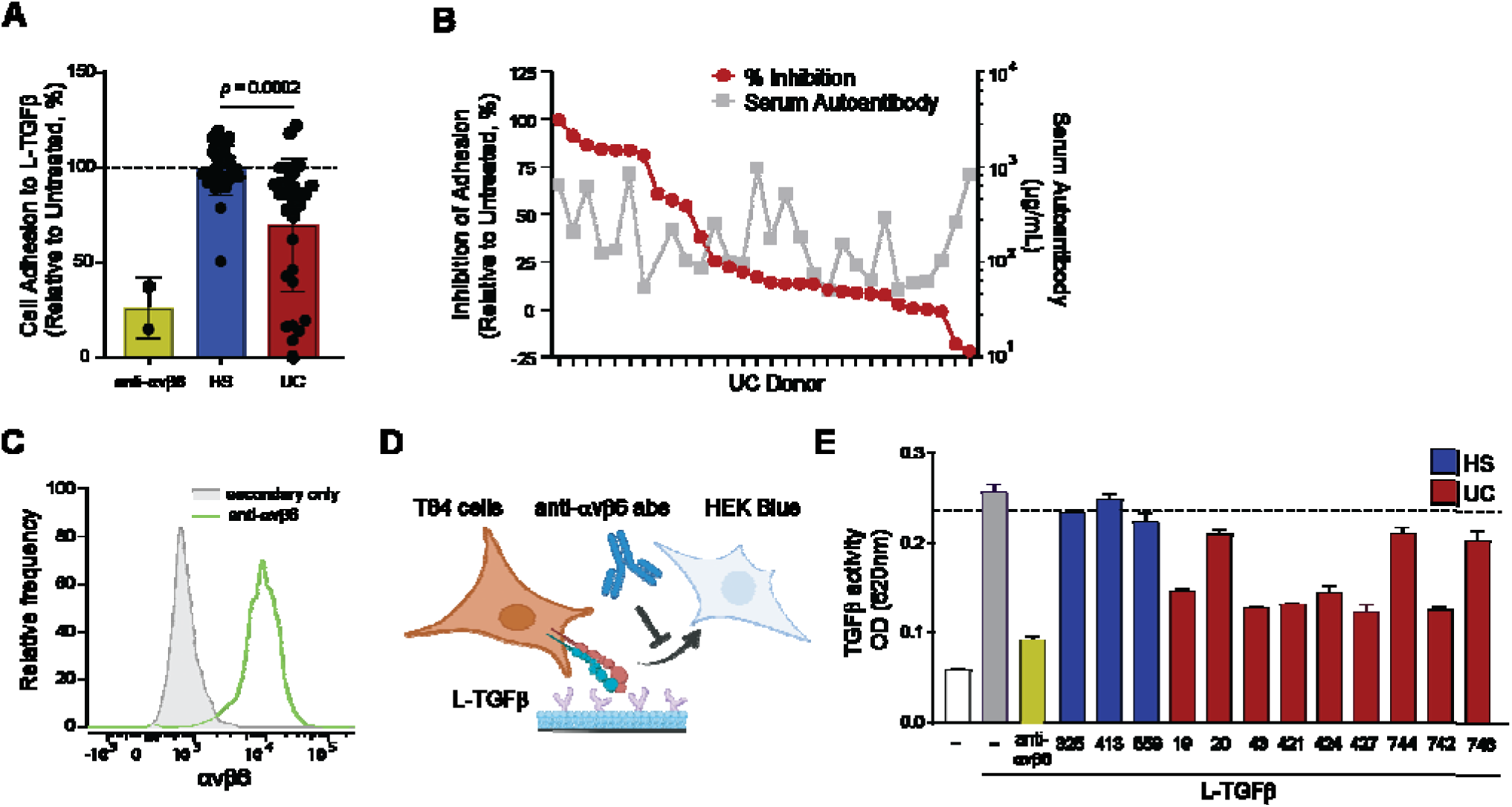
Functional effects of UC-derived anti-αvβ6 autoantibodies on αvβ6-mediated adhesion and activation of TGFβ. **(A)** Adhesion of HT-29 cells to latent TGFβ (L-TGFβ) in the presence of anti-αvβ6 blocking antibody 3G9, HS serum, or UC serum, normalized to untreated controls. **(B)** Inhibition of cell adhesion to L-TGFβ by individual donor UC serum (left axis) plotted against corresponding serum anti-αvβ6 IgG levels (right axis). **(C)** Representative flow cytometry histograms showing surface expression of αvβ6 on T84 cells stained with 3G9 (green) or secondary-only control (gray). **(D)** Schematic of the in vitro assay used to measure αvβ6-dependent activation of L-TGFβ by T84 cells and signaling to HEK-Blue^TM^ TGFβ reporter cells. **(E)** αvβ6-dependent activation of latent TGFβ by T84 cells measured by HEK-Blue reporter activity (OD620); dotted line indicates mean reporter activity across HS controls. Data represent mean ± SD and statistical significance determined using unpaired two-tailed t test with Welch’s correction **(A)**, individual data points **(B)** or mean ± SEM **(E)**

To test whether UC autoantibodies inhibit activation of TGFβ, we co-cultured T84 colonic epithelial cells, which express αvβ6 (**Fig 2C**), with a TGFβ signaling reporter cell line in the presence of L-TGFβ (**Fig 2D**). We confirmed that T84 cells were able to activate L-TGFβ for signaling to co-cultured TGFβ reporter cells, and this was inhibited by the monoclonal antibody to αvβ6, 3G9 (**Fig 2E).** We next selected UC sera that effectively blocked L-TGFβ binding, and purified IgG to exclude confounding effects of TGFβ present in serum. IgG from individuals with UC inhibited activation of TGFβ while IgG from HS that were negative for anti-αvβ6 antibodies had no effect **(Fig 2E**). To exclude the possibility that the IgG-mediated inhibition of TGFβ activation by colonic epithelial cells was not mediated through blockade of αvβ6, we generated TGFβ reporter cells expressing the β6 subunit (β6 HEK-Blues) **(Supp Fig 2C**).

Wildtype HEK-Blue cells express αv integrin but not β6, and are unable to activate L-TGFβ, whereas β6 HEK-Blues gain the ability to activate L-TGFβ (**Supp Fig 2D,E**). L-TGFβ activation by β6 HEK-Blues was equally blocked by 3G9, or by IgG from a UC donor that showed blocking activity in **Fig 2E** (**Supp Fig 2F**). Together, these data show that UC autoantibodies inhibit αvβ6-dependent binding of colonic epithelial cells to L-TGFβ and activation of L-TGFβ for signaling.

### Inhibition of **α**v**β**6-mediated TGF**β** activation disrupts epithelial cell homeostasis

Increasing evidence indicates that UC is associated with intrinsic remodeling of the intestinal epithelium. Analyses of human colonic tissue have revealed widespread alterations in epithelial cell states, differentiation trajectories, and stress and repair–associated transcriptional programs in UC (*40–42*). Given the established role of TGFβ signaling in regulating epithelial proliferation, differentiation, and repair in the intestine (*28, 29*), we examined the transcriptional consequences of epithelial TGFβ inhibition downstream of αvβ6 blockade.

T84 cells were differentiated into mature colonic epithelial monolayers and treated daily for 5 days with the anti-αvβ6 monoclonal antibody 3G9, a TGFβ neutralizing antibody, active TGFβ, or media-alone, followed by transcriptional analysis by RNA-Seq. We identified 62 differentially expressed genes (DEGs) following anti-αvβ6-treatment relative to untreated controls (**Fig 3A**), and 318 DEGs in response to active TGFβ **(Fig 3B**). Treatment with a TGFβ neutralizing antibody also caused gene expression changes in T84 cells (**Supp Fig 3A**), and although these were smaller in magnitude than with αvβ6 blockade, we identified a shared set of anti-αvβ6 and anti-TGFβ-responsive genes whose directionality was generally conserved across both treatments (**Fig 3C**). The majority of these shared DEGs were regulated in the opposite direction by treatment with TGFβ, indicating that they are true TGFβ-responsive genes. A subset of transcripts was more strongly modulated by anti-αvβ6 than by TGFβ neutralization. Notably, αvβ6 inhibition increased expression of the transport genes *AQP8* and *SLC26A3*, while decreasing *SLC30A2*, indicating shifts in epithelial fluid and electrolyte handling (53–55). In parallel, multiple metallothionein family members (MT1 and MT2), which bind intracellular zinc and have been linked to protection against intestinal inflammation, were downregulated (56–61). Given that *SLC30A2* encodes a zinc exporter and MTs regulate intracellular zinc buffering, these coordinated changes suggest altered epithelial zinc homeostasis following αvβ6 blockade. Together, these findings indicate that αvβ6 inhibition primarily disrupts TGFβ-responsive epithelial programs, while exerting additional selective effects on zinc-handling and transport pathways.

**Fig. 3.**
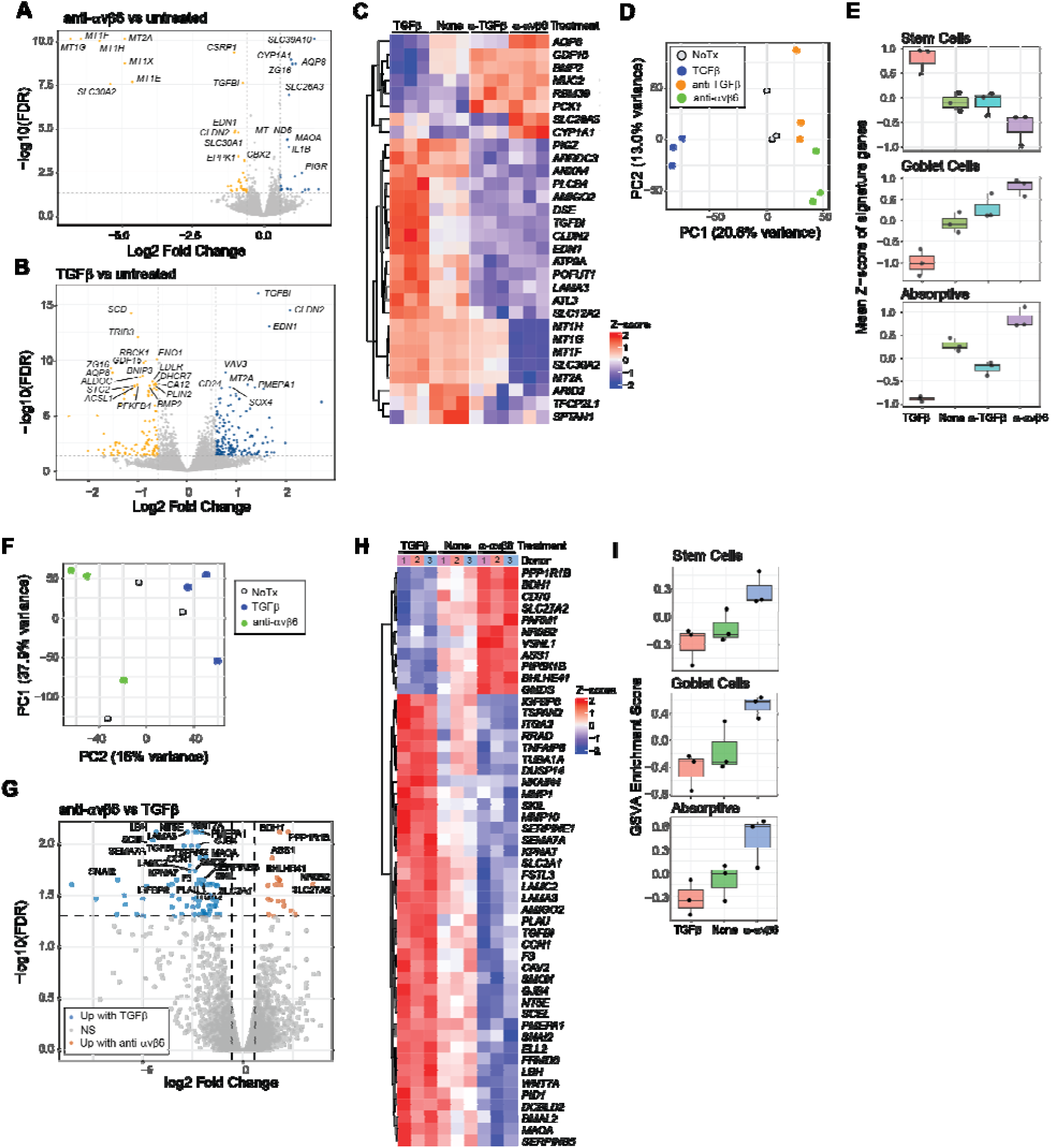
Transcriptional profiling of human IECs treated with anti-αvβ6 or TGFβ. **(A-B)** Volcano plot of differentially expressed genes (DEGs) for **(A)** 3G9-treated vs untreated T84 cells or **(B)** TGFβ-treated vs untreated T84 cells. **(C)** Heatmap of genes commonly regulated by 3G9 and anti-TGFβ relative to untreated controls. **(D)** Principal component analysis (PCA) of T84 cell transcriptomes. **(E)** Epithelial subset signature scores in T84 cells based on curated marker gene sets. **(F)** PCA of primary IEC transcriptomes. **(G)** Volcano plot of DEGs for 3G9- vs TGFβ-treated primary IECs. **(H)** Heatmap of DEGs between 3G9- and TGFβ-treated primary IECs. **(I)** Epithelial subset program enrichment in primary IECs quantified by gene set variation analysis (GSVA). Dotted lines in volcano plots indicate fold-change and false discovery rate (FDR) thresholds.

Principal Component Analysis (PCA) clearly separated TGFβ-treated, untreated, and anti-TGFβ treated cells, with cells treated with 3G9 clustering closest to anti-TGFβ-treated cells **(Fig 3D)**. To further define the broader transcriptional programs associated with TGFβ signaling, we next examined gene ontology (GO) enrichment analysis of genes contributing positively to PC1 (corresponding to TGFβ signaling). We found that PC1 genes enriched for processes related to translational control, differentiation, regulation of growth, and cellular stress responses (**Supp Fig 3B**). This profile suggests that TGFβ signaling coordinates upregulation of stress-adaptive and maintenance pathways which are downregulated following αvβ6 or TGFβ blockade in differentiated colonocytes.

Genes negatively associated with PC1 (corresponding to inhibition of TGFβ signaling) did not demonstrate significant GO term enrichment, prompting deeper examination of the individual DEGs shared between anti-αvβ6 and anti-TGFβ treatment (**Fig 3C**). Among these were several downregulated genes involved in epithelial adhesion and matrix organization, including *TGFBI*, *LAMA3*, *DSE*, *AMIGO2*, and *ARRDC3* (*43–46*), and the tight junction component *CLDN2* (*46*). These findings indicate that inhibition of αvβ6-mediated TGFβ activation alters structural and barrier-associated gene programs. Additionally, secretory goblet cell-associated genes were also affected. Blockade of αvβ6 or TGFβ signaling increased expression of *MUC2*, *BMP2*, and *GDF15* and reduced expression of the Notch-associated regulator *POFUT1*, consistent with enhancement of goblet cell programs (*47–49*).

The induction of goblet cell-associated genes together with marked changes in epithelial fluid and electrolyte transport genes raised the possibility that loss of αvβ6 function impacts epithelial differentiation programs of absorptive and goblet IEC lineages. To determine whether transcriptional changes reflected broader shifts in epithelial cell state, we assessed lineage-associated epithelial programs using curated marker gene signatures from human colon scRNA-Seq studies (*6, 40*). For each IEC subset, we calculated per-sample signature scores based on the mean Z-score across marker genes and treatment conditions. TGFβ treatment suppressed absorptive and goblet cell expression (**Fig. 3E, Supp Fig 3C)**, consistent with a role for TGFβ signaling in restraining epithelial differentiation. In contrast, anti-αvβ6 led to an induction of both absorptive and goblet cell signatures, with more pronounced effects than those observed following pan-TGFβ blockade, while stem cell gene signatures showed opposing regulation.

### **α**v**β**6-dependent TGF**β** signaling maintains differentiation programs in primary IECs

While differentiated T84 cells serve as a well-controlled system to study effects on terminally differentiated IECs, they are limited in their ability to model early differentiation dynamics and lineage commitment. We therefore turned to an *in vitro* culture system using primary human IECs derived from colon biopsies of healthy individuals which supports maintenance of intestinal stem cells and their differentiated progeny (*50*). Primary IECs were treated once with anti-αvβ6, active TGFβ, or media-alone, and gene expression measured by RNA-Seq after 4 days in culture. PCA revealed that the dominant source of variance reflected donor differences (PC1), as expected for primary human samples, while PC2 captured TGFβ-dependent differences (**Fig 3F)**. As observed for T84 cells, TGFβ-treated and anti-αvβ6 treated samples separated along this component with untreated samples positioned in between, indicating that primary IECs also undergo basal TGFβ activation and signaling that is inhibited by αvβ6 blockade.

Genes associated positively with PC2 (TGFβ response) showed enrichment for wound healing and coagulation, cell-matrix adhesion, and multiple Wnt signaling modules, consistent with known effects of TGFβ signaling (**Supp Fig 3D**). Notably, enriched Wnt-related terms included both canonical and non-canonical Wnt pathways as well as developmental programs linked to epithelial and connective tissue differentiation, consistent with known crosstalk between TGFβ and Wnt signaling in regulating epithelial fate decisions (*51, 52*).

Pairwise comparisons between 3G9 and TGFβ-treated IECs identified 90 DEGs, with the majority displaying intermediate expression in untreated cells, confirming that anti-αvβ6 inhibits TGFβ signaling in these cultures (**Fig 3G**). DEGs included genes reported in canonical TGFβ transcriptional programs that were downregulated by αvβ6 blockade, including *SERPINE1*, *TGFBI*, *PMEPA1*, *CCN1*, *SKIL*, and *SNAI2* (*43, 53*). αvβ6 inhibition also decreased expression of multiple genes involved in extracellular matrix organization and adhesion including *LAMA3*, *LAMC2*, *ITGA2*, *TSPAN2*, *PLAU*, and *MMP10* (**Fig 3H**) (*53, 54*). αvβ6-mediated TGFβ signaling in primary human IECs therefore promotes a coordinated program in developing epithelial cells encompassing canonical TGFβ target gene expression, extracellular matrix remodeling, tissue repair pathways, and Wnt-signaling dependent differentiation.

We next analyzed expression of the same lineage-associated gene signatures used in our differentiated T84 cell dataset (**Supp Fig 3E**). Anti-αvβ6 promoted expression of genes associated with absorptive and goblet cell programs, whereas TGFβ treatment showed the opposite pattern (**Fig. 3I**). Interestingly, anti-αvβ6 caused increased expression of some stem cell-associated genes, the opposite effect observed in T84 cells, although the effects were weaker in primary IECs. This likely reflects context-dependent effects of TGFβ signaling which can differentially regulate stem and progenitor populations versus terminally differentiated epithelial cells. Collectively these results indicate that intestinal epithelial cells rely on αvβ6-mediated TGFβ activation and signaling in culture, which constrains epithelial plasticity in primary IECs and regulates differentiation trajectories. Inhibition of αvβ6 promotes goblet and absorptive epithelial cell programs in both early developing IECs and fully differentiated cells.

### Spatial transcriptomics reveal cellular remodeling in **α**v-villin mouse colon

*In vivo*, epithelial cell state is influenced by multiple cell intrinsic and extrinsic factors, including tissue location, local immune and stromal cells, and microbes in the gut lumen. Furthermore, intestinal epithelial cells are known to direct differentiation of gut resident immune cells, in part through TGFβ activation (*55*). To understand how inhibition of epithelial αvβ6 function affected epithelial and immune homeostasis, we generated a mouse model in which αv integrins were deleted from intestinal epithelial cells, termed αv-villin mice.

αv-villin mice exhibited no developmental defects or evidence of intestinal inflammation (**Supp Fig 4A,B).** This is consistent with the phenotype of total *Itgb6* knockouts, which also show no intestinal inflammation but do exhibit some spontaneous skin inflammation (*56*). To more deeply probe changes in the intestinal epithelium, we analyzed ‘swiss roll’ preparations of colon from 12-week-old αv-villin and littermate control mice using the Visium HD spatial transcriptomics platform **(Fig 4A**).

**Fig. 4.**
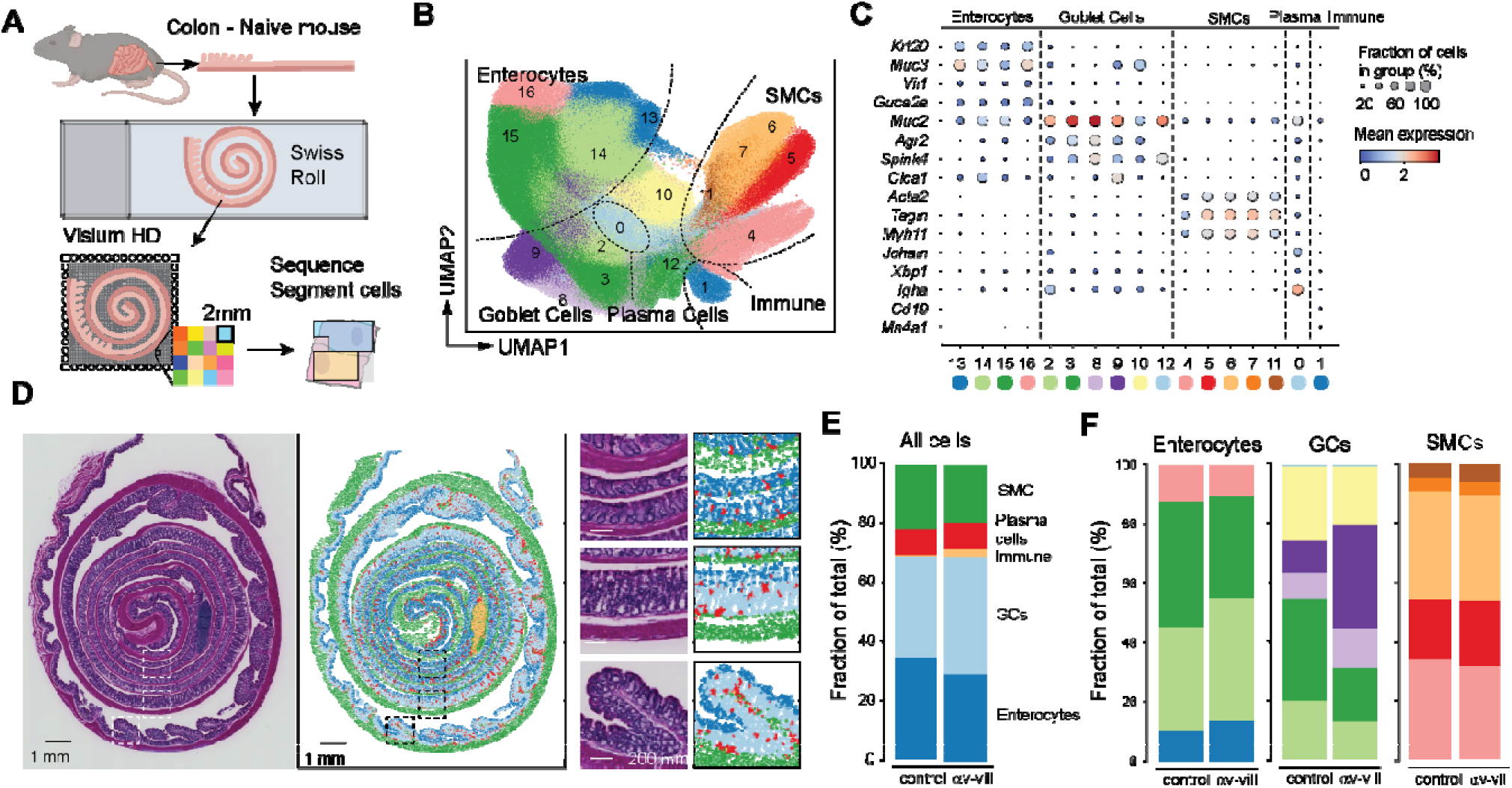
Spatial transcriptomics analysis of colon from αv-villin and control mice. **(A)** Schematic of experimental design using ‘swiss roll’ sections of colons from naïve αv-villin and control mice analyzed by Visium HD. **(B)** UMAP projection of leiden clusters of all cell types identified in 2 control and 2 αv-villin mice. **(C)** Dot plot with relative expression of selected genes in cell clusters. **(D)** Projection of major cell type clusters onto swiss roll histological sections (right) with H&E stain (left). Images captured at 4X, scale bars 1mm and 200μm. Small images show magnification of regions indicated by dashed lines. Images are from one representative αv-villin mouse. **(E-F)** Relative proportions of cell clusters in control and αv-villin mice. Left panel shows all clusters together grouped by cell type; right panel shows relative proportions of clusters within enterocytes, GCs and SMCs.

We identified a total of 727,091 cells across all samples. After clustering, cells were separated into 5 major cell types (smooth muscle cells [SMCs], plasma B cells, other immune cells, and 2 major types of epithelial cells, enterocytes and goblet cells [GCs**])(Fig 4B,C**). These cell types mapped to expected regions of the colon **(Fig 4D**). With the exception of plasma B cells, which were found throughout the lamina propria, other immune cells were largely confined to large immune follicles. In this experiment, follicles were most prominent in αv-villin mice but were seen in other colon sections from control mice, and therefore represent variation in tissue orientation rather than underlying differences due to genotype. Other cell populations including neurons, endothelial cells, and lamina propria T cells and dendritic cells (DCs) could not be unambiguously identified due to their small size compared with the resolution of Visium HD (2 μm), and high background of highly expressed epithelial transcripts such as *Muc2.* Initial analyses of the 5 major cell types identified differences in the GC compartment of αv-villin colon, with an increase in the fraction of total GCs and changes in the relative proportion of individual GC clusters relative to controls (**Fig 4E,F**).

### Loss of epithelial **α**v causes expansion of canonical goblet cells in the mid colon

To better define the changes observed in the GC compartment in αv-villin mice, GCs and enterocytes were separated based on relative expression of Muc2 and Muc3 and re-clustered. Subclusters showed major differences in distribution along the colon, with clear GC and enterocyte populations restricted to proximal, mid and distal regions, and distinct positioning along the crypt-lumen axis (**Fig 5A-D)**, in agreement with other scRNA-Seq and spatial transcriptomics studies (41, 73–77). Comparison of cluster frequencies between genotypes revealed that several GC subsets were increased in αv-villin mice (**Fig 5E**, left). When projected onto H&E images, the expanded clusters appeared to be localized to the mid colon (**Supp Fig 5A,B)**, and this was confirmed by digital ‘unrolling’ of colon images (**Fig 5E**, right). By contrast, enterocyte proportions and spatial distributions were largely unchanged by genotype (**Supp Fig 5C**).

**Fig. 5.**
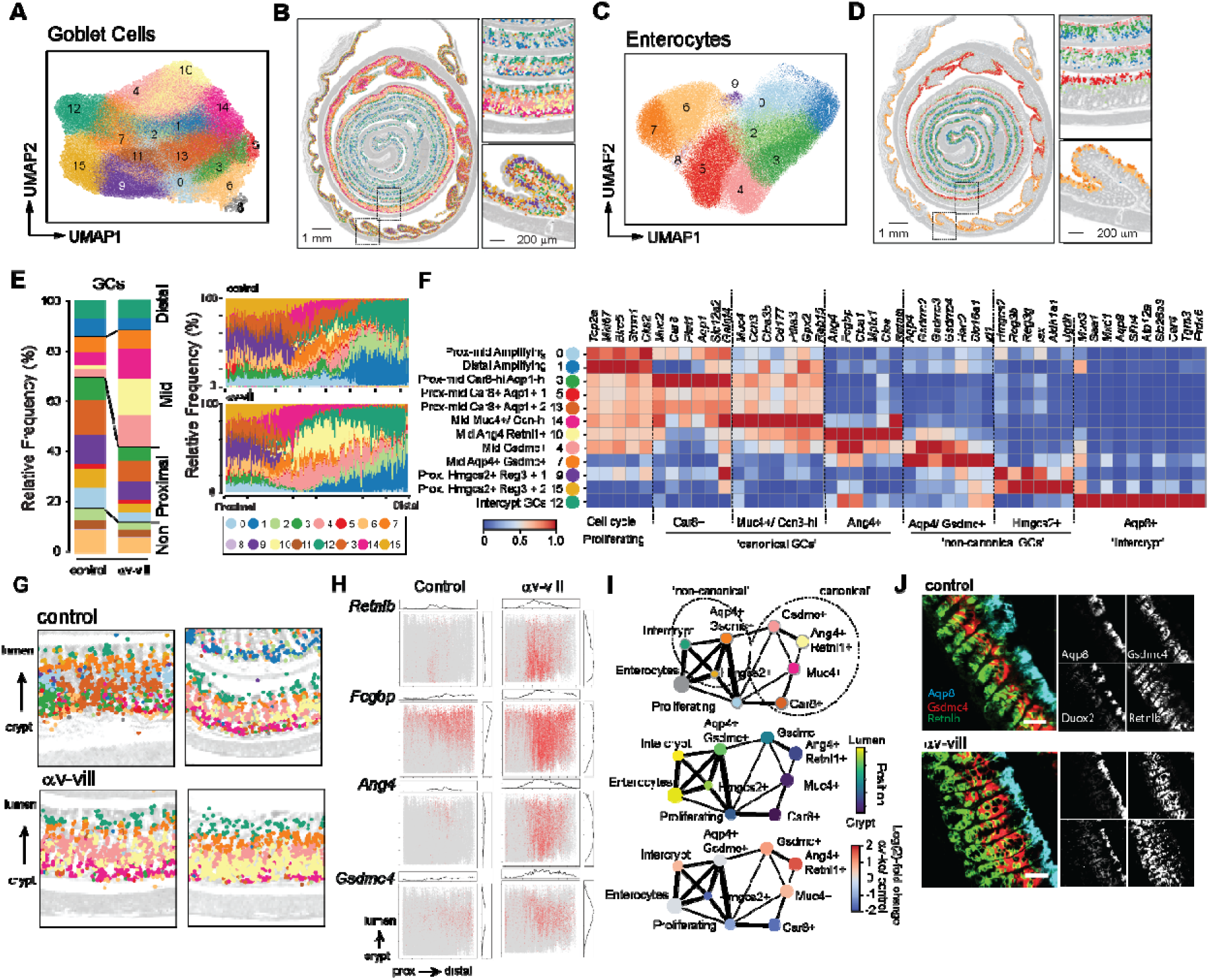
Characterization of expanded goblet cells in αv-villin mice. **(A-D)** Leiden clusters of GCs **(A)** or enterocytes **(C)** projected by UMAP for all mice, or spatially **(B, D)** for a representative αv-villin mouse. Small images show magnification of regions indicated by dashed lines. **(E)** Relative proportions of GC clusters in control and αv-villin mice. Left panel shows relative frequency across the whole colon, right panels show frequencies for sequential regions along the length of the colon. **(F)** Heat map showing Z-score scaling of log-normalized goblet cell signature gene expression across GC clusters. **(G)** Spatial projection of GC clusters, colored as in **(E)**, over H&E image (grayscale) for mid colon. Each image is from an individual mouse. **(H)** Trajectory analysis of enterocytes and GCs from the mid colon (2 control and 2 αv-villin mice were combined). Plots show cell subsets colored by cluster identify as in **(F)** (top), by median position in the crypt-lumen axis (middle), and by relative abundance in αv-villin versus controls (bottom). **(I)** Expression of indicated genes in cells plotted on unrolled and aligned axis (colon length and crypt to lumen). (J) mRNA *in situ* hybridization of *Aqp8, Gsdmc4, Retnlb* and *Duox2* in mid colon of αv-villin and control mice. 10x objective, scale bar 50μm.

Colonic GCs comprise multiple functionally distinct subsets that differ by location and gene expression profile. To define the identities of GC populations altered in αv-villin mice, we integrated differential expression analysis between GC subclusters with curated marker gene signatures from published mouse and human colon datasets (41, 42, 73–78). Twelve clusters could be classified into seven major GC subsets (**Fig 5F**), while four subsets included contaminating non-epithelial transcripts that were excluded from further analysis. Assigned GC subsets included *(i)* recently proliferating cells undergoing early GC differentiation (clusters 0 and 1) characterized by high levels of *Mki67* and *Top2a* located in the low to mid crypt; *(ii)* three populations of ‘canonical’ GCs, defined by expression of *Car8* and *Aqp1* (clusters 3, 5 and 13), *Ccn3* and *Muc4* (cluster 14), or *Ang4, Fcgbp* and *Clca1* (cluster 10) (*57*); *(iii)* two populations of ‘non-canonical’ GCs expressing intermediate levels of *Muc3* and either *Hmgcs2* and *Reg3b/g* (clusters 9 and 15) or *Aqp4, Gsdmc4* and *Hao2* (cluster 7) (*57*); and *(iv)* a population uniquely expressing *Mxd1, Slfn1* and *Aqp8* (Cluster 12), which was predominantly present in the distal colon and restricted to the luminal surface, consistent with intercrypt GCs (*41*).

The 3 expanded mid colon GC populations in αv-villin mice (4, 10 and 14) had distinct but overlapping gene expression patterns and were enriched for antimicrobial and inflammation-associated genes upregulated in infection, such as *Retnlb, Ang4* and *Mptx1* (*58–60*), and *Gpx2* and *Pla2g4c,* which encode proteins involved in superoxide production and oxidative stress. Cluster 14 also expressed *Muc4, Ccn3* and *Pdia3*, and represented a subpopulation of GCs found in the lower crypt that are specialized for acidic mucin production (*61*). Cluster 10 expressed high levels of *Ang4,* and closely resembled GCs that develop in mice during bacterial colonization (*57*). Cluster 4 shared expression of several signature genes with the *Ang4+* GCs (cluster 10) but also expressed high levels of genes encoding Gsdmc.

These expanded clusters were ordered along the crypt-lumen axis with cluster 14 closest to the crypt, followed by cluster 10 and then cluster 4 in the mid-crypt region (**Fig 5G; Supp Fig 5D)**. Four ‘signature’ GC genes that showed overlapping expression for each subset, *Retnlb, Ang4, Gsdmc* and *Fcgbp,* showed similar patterns, with *Retnlb* and *Ang4* expressed closer to the crypt, *Gsdmc4* in the mid crypt, and *Fcgbp* expanded throughout the crypt–lumen axis in αv-villin mice (**Fig 5H**). We speculated this may reflect the maturation of GCs as they migrate up the crypt, particularly as all three subsets expressed genes consistent with canonical GCs which are proposed to arise along a distinct developmental trajectory to non-canonical GCs (*41*). In support of this model, we identified 2 major trajectories in the mid colon which separated the canonical GCs from the non-canonical subsets, which were more closely related to enterocytes (**Fig 5I**). All 3 expanded subsets laid within the canonical trajectory and were ordered consistently with their positions within the crypt. The expansion of these subsets accompanied by fewer proliferative GCs in the mid colon in αv-villin mice suggest that loss of αv biases GC differentiation towards more mature states, consistent with the skewing towards GCs observed in human IECs (**Fig 3E,I**), and that this predominantly affects the canonical GC pathway.

To confirm the locations of these GC populations, we assessed sections from additional control and αv-villin mice for *in situ* expression of 4 genes: *Retnlb*, expressed by the expanded canonical GC subset; *Gsdmc4*, expressed by both expanded canonical and non-canonical GCs; and *Aqp8* and *Duoxa2*, which are expressed in intercrypt cells. As seen by spatial transcriptomics, *Retnl1b* was expressed in GCs close to the crypt and *Gsdmc4* in cells midway along the crypt. Both populations were expanded in αv-villin mice but had largely non-overlapping patterns of gene expression with few *Retnlb* and *Gsdmc4* double positive cells (**Fig 5J**). By contrast, *Aqp8* and *Duoxa2* were expressed in intercrypt cells and did not show major changes in the pattern of expression in αv-villin mice. Together, these data reveal that disruption of epithelial αv causes expansion of subsets of GCs involved in anti-microbial response and inflammation, and that effects were primarily localized to crypt cells in the mid-colon.

### Epithelial **α**v regulates intraepithelial immune cell residency

The epithelium exists in close proximity to immune cells, including specialized intra-epithelial lymphocytes (IELs) and populations of macrophages and dendritic cells (DCs). The differentiation, retention, and regulation of these immune cells rely on signaling from TGFβ that is activated locally by epithelial cells, and deletion of β6 from epithelial cells in the skin and small intestine results in loss of IELs (*55*). We therefore investigated whether deletion of αv affected immune cell populations in the colon. αv-villin mice had similar total numbers of colonic IELs to control mice (**Fig 6A,B)** but had a selective loss of ‘conventional’ TCRαβ^+^ CD8αβ^+^ IELs (**Fig 6C**). Furthermore, expression of CD103 (αEβ7 integrin), a marker of tissue resident lymphocytes that contributes to lymphocyte retention through binding to E-cadherin on epithelial cells (*62, 63*), was completely lost in αv-villin mice **(Fig 6D**). Similar loss of CD103 was observed in the other major populations of IELs, TCRαβ^+^ CD8αα^+^ and TCRγδ^+^ cells (**Fig 6E,F**). In T cells, CD103 is strongly responsive to TGFβ, and loss of expression on IELs therefore supports a role for epithelial αv in local activation of TGFβ for signaling *in trans* to interacting lymphocytes.

**Fig. 6.**
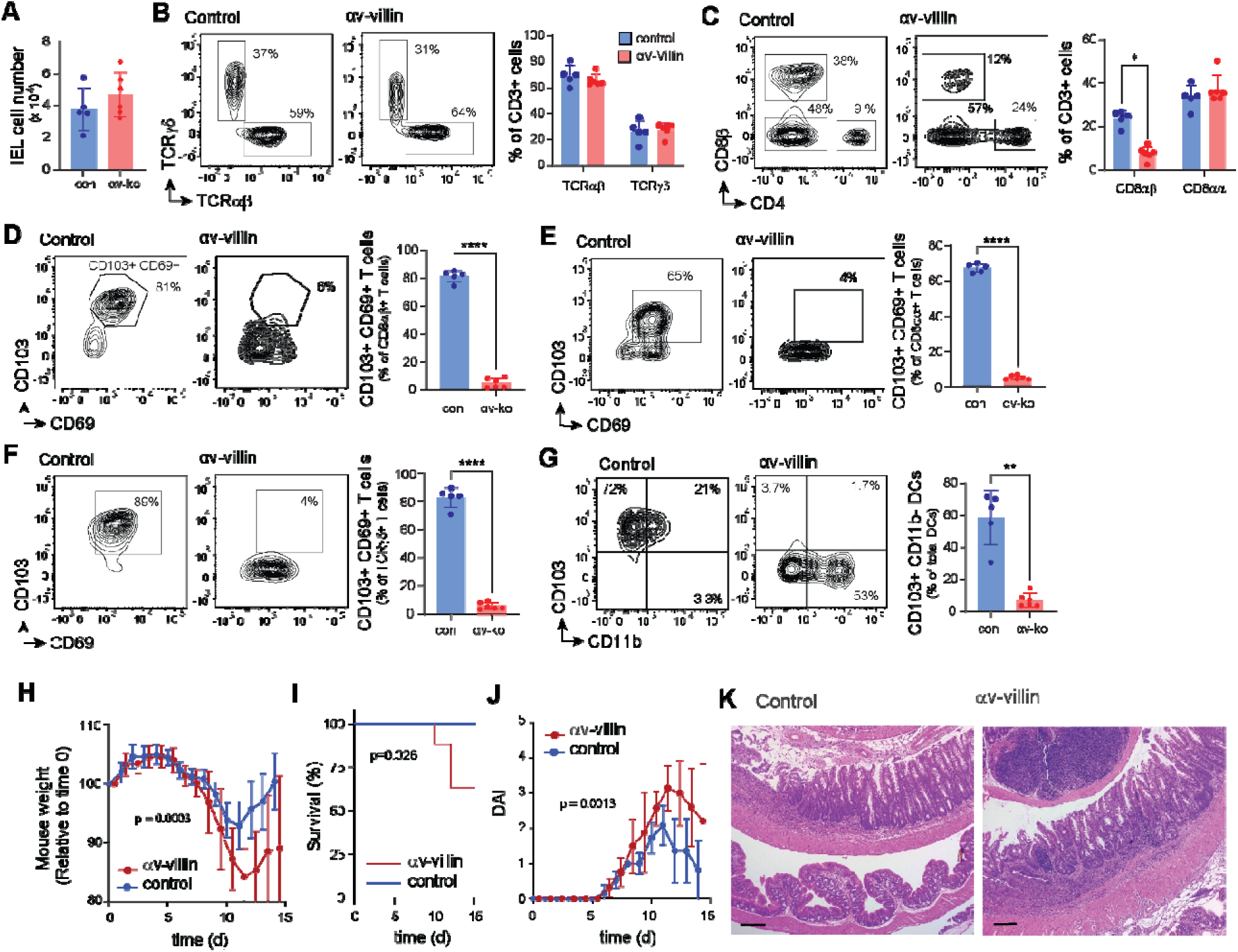
Intestinal immune cell composition and DSS-colitis responses in αv-villin mice. **(A)** Total number of intraepithelial lymphocytes (IELs; Live/Dead^-^CD45^+^ cells) from WT and αv-villin mice. **(B-G)** Representative plots and frequencies of TCRαß and TCRγδ IEL of CD3^+^ T cells **(B)**, CD8αß and CD8αα IEL of CD3^+^ T cells **(C)**, CD103^+^CD69^+^ IEL of CD8αß^+^ T cells **(D)**. Representative plots and frequencies of CD103^+^CD69^+^ IEL of CD8αα^+^ T cells **(E)**, CD103^+^CD69^+^ IEL of TCRγδ^+^T cells **(F)**, CD103^+^CD11b^-^ DCs of MHCII^+^CD11c^+^ DCs **(G)**. **(H)** Body weight, **(I)** Kaplan Meier survival curve, **(J)** disease activity index (DAI), and **(K)** representative colonic H&E (4X, scale 100 μm) at day 14 following 2% DSS. Flow cytometry data are shown as mean ± SD and are representative of three independent experiments with n = 5 (WT) and n = 6 (αv-villin) mice per group. Statistical significance was determined using unpaired t tests with Welch’s correction **(A, D-G)** or two-stage step-up FDR correction **(B-C)**. Body weight and DAI (mean ± SEM) were analyzed using mixed-effects models (REML; p = 0.0003 and p = 0.0013, respectively). Survival was analyzed using Kaplan–Meier curves and compared using the log-rank (Mantel–Cox) test (p=0.028). DAI were analyzed using a mixed-effects model (REML; p = 0.0013). * = P < 0.05; ** = P < 0.01; *** = P < 0.001; **** = P < 0.0001

We also analyzed CD11c^+^ DCs which were isolated in the colon IEL fraction. In control mice, these IEL DCs include CD103^+^ CD11b^-^ cDC1s, which are involved in modulation of epithelial inflammatory responses and generation of Tregs, and CD103^+^ CD11b^+^ cDC2s, which are associated with Th17 responses (*64–67*). CD103 expression was lost in IEL DCs from αv-villin mice and a relative increase in CD11b^+^ DC2s (**Fig 6G**).

In contrast with the effects on IELs, we observed no major differences in the number of colon LP immune cells, or in activated CD4 and Treg T cell numbers in αv-villin mice (**Supp Fig 6A-C).** We also did not see significant changes in T cells or DCs in the mesenteric lymph node (MLN) of αv-villin mice compared with controls (**Supp Fig 6D-I**). Hence epithelial αv plays a non-redundant role for generation of CD103^+^ T cells and DCs resident in the colonic epithelium.

### Deletion of epithelial **α**v exacerbates colitis in mice

Despite changes in the intestinal epithelium and resident immune cells, αv-villin mice did not develop spontaneous inflammation. To determine whether loss of epithelial αv affected the development of colitis, mice were treated with Dextran Sodium Sulphate (DSS). αv-villin mice developed more severe disease than littermate controls, showing increased weight loss, mortality, and disease score **(Fig 6 H-J**). This was particularly apparent during days 8 to 14, when mice returned to normal drinking water. When harvested after 14 days, the majority of control mice had recovered their weight loss whereas αv-villin mice had persistent weight loss, disease activity, and more severe intestinal inflammation (**Fig 6 H-K**).

### **α**v**β**6 autoantibodies block epithelial TGF**β** activation and exacerbate colitis *in vivo*

Taken together, these data demonstrate that deletion of αv integrins from the intestinal epithelium disrupts the colon GC and immune cell compartments and exacerbates experimental colitis. However, it remains unclear whether development of autoantibodies in fully developed mice could recapitulate the effects of genetic deletion. To better model the development of UC associated αvβ6 autoantibodies, we established a novel murine autoimmune model. C57BL/6 mice were immunized with recombinant human αvβ6 protein emulsified in Complete Freund’s Adjuvant (CFA). Reflecting the 89.5% homology between human and mouse αvβ6, this approach elicited high serum titers of antibodies that recognized both human and mouse αvβ6 while no anti-αvβ6 activity was seen in control mice that received adjuvant alone **(Fig. 7A)**. Sera from mice immunized with αvβ6 were function blocking, inhibiting cellular adhesion of HT-29 cells to L-TGFβ in a dose-dependent manner **(Fig. 7B)**. Immunized mice did not develop spontaneous intestinal inflammation, even up to 12 weeks after immunization (**Fig 7C,D**). We noted that immunized mice did develop hair loss and skin lesions at sites of immunization, and epithelial hyperplasia around ear tags (**Supp Fig 7A,B**), similar to phenotypes seen in *Itgb6^-/-^*mice (*68*), indicating that αvβ6 autoantibodies inhibit function *in vivo*. As observed in αv-villin mice, persistence of epithelium-resident immune cells was impaired in αvβ6-immunized mice, with reduced numbers of CD8αβ ^+^ IELs and loss of CD103 expression **(Fig 7E,F**). Finally, in the DSS colitis model, αvβ6-immunized mice had greater weight loss and lower survival than adjuvant-alone controls (**Fig 7G,H**). After harvest at day 14, αvβ6-immunized mice had shorter colon lengths and more severe inflammation than controls **(Fig. 7I,J)**.

**Fig. 7.**
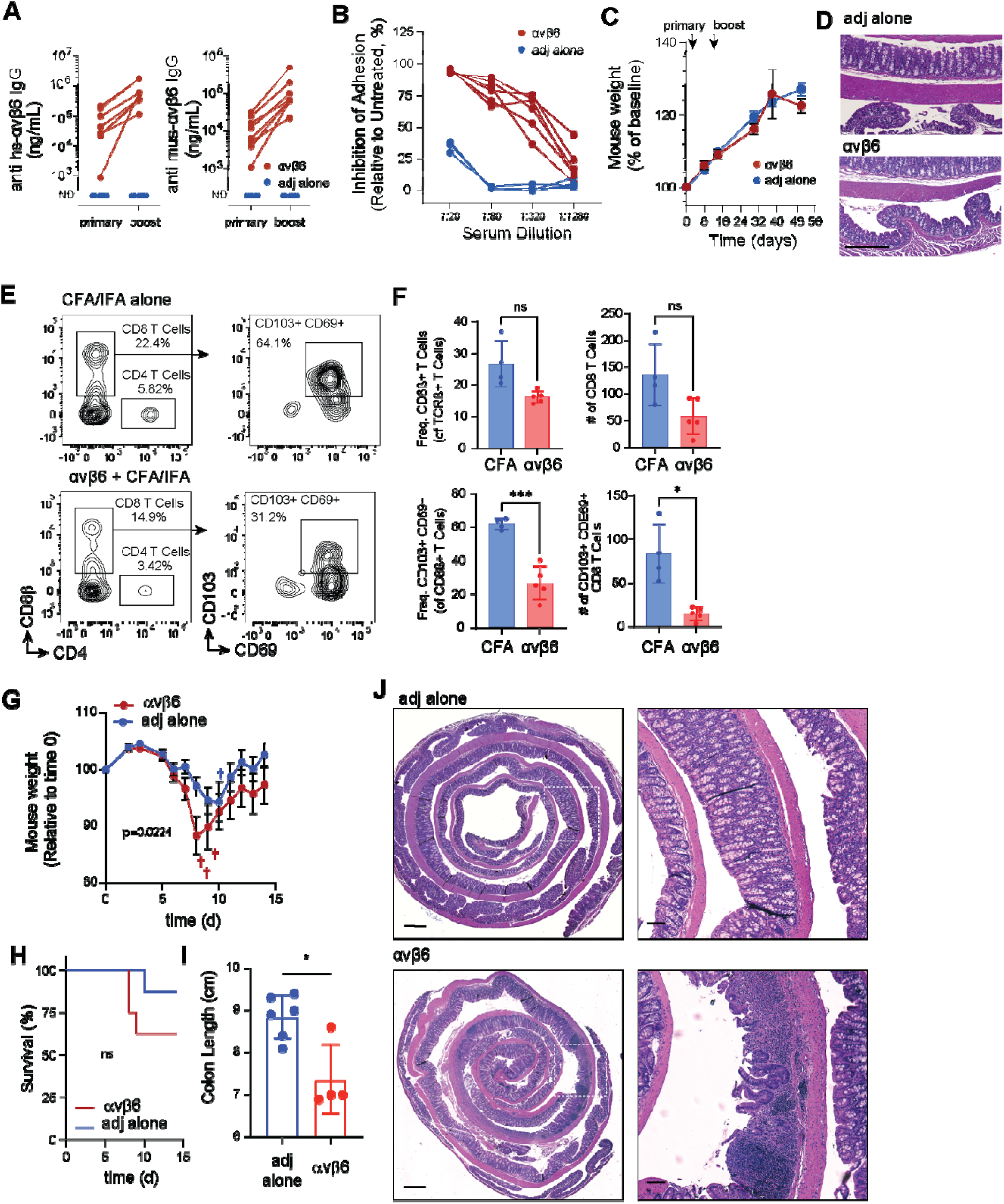
Induction of anti-αvβ6 autoantibodies and responses to DSS-colitis. **(A)** Serum anti-human (left) and anti-mouse (right) αvβ6 IgG in αvβ6-immunized and adjuvant-only (n=8 per group) mice quantified by ELISA following primary immunization and boost. **(B)** Inhibition of αvβ6-dependent adhesion to L-TGFβ by serial dilutions of serum from αvβ6-immunized (n=6) or adjuvant-only (n=4) mice. **(C)** Body weight following primary immunization and boost. **(D)** Representative colonic H&E at baseline. **(E, F)** Representative flow cytometry plots and frequencies of CD8αß^+^ T cells among TCRß^+^ T cells **(E)** and CD103^+^CD69^+^ T cells among CD8ß^+^ T cells **(F)** from IELs. **(H-K)** Following DSS-induced colitis, body weight **(H)**, survival **(I)**, colon length **(J)**, and representative H&E-stained colon sections at day 14 post-DSS **(K)**. Body weight data (mean ± SEM) were analyzed using a mixed-effects model (REML; p = 0.0224). Survival was analyzed using Kaplan–Meier curves and compared by the log-rank (Mantel–Cox) test (p=0.22). Flow cytometry data are shown as mean ± SD and are representative of a single experiment (of two independent experiments) conducted with n = 4 adjuvant-only and n = 5 αvβ6-immunized mice per group. Statistical significance was determined using multiple unpaired t tests with Welch’s correction **(F, H, J)** or log-rank (Mantel–Cox) test **(I)**. *P < 0.05; **P < 0.01; ***P < 0.001

## Discussion

UC is characterized by epithelial barrier dysfunction and sustained mucosal inflammation; however, the mechanisms that initiate disease remain incompletely understood. αvβ6 autoantibodies have emerged as specific biomarkers of UC that can precede clinical diagnosis by up to 10 years (*9–15*). Here, we show that these autoantibodies are functional, blocking αvβ6-dependent activation of latent TGFβ in the intestinal epithelium. This inhibition altered epithelial differentiation programs, disrupted epithelial–immune crosstalk, and increased susceptibility to colitis *in vivo* in mouse models.

Across human and mouse systems, deletion or inhibition of αvβ6 revealed that epithelial cells rely on ongoing αvβ6-mediated TGFβ signaling to maintain coordinated differentiation and barrier-associated gene programs. In human IECs, αvβ6 blockade altered expression of genes involved in epithelial transport, tight junction regulation, adhesion, and extracellular matrix organization, including *AQP8* and metallothionein genes which are implicated in epithelial homeostasis and dysregulated in UC mucosa (*69–75*). αvβ6 blockade enhanced GC lineage-associated gene signatures *in vitro*, and epithelial αv deletion *in vivo* led to expansion of canonical goblet cell populations in the mid colon. Notably GC-associated genes induced by αvβ6 blockade in T84 cells (*MUC2*, *RETNLB*, *FCGBP*, *CLCA1*) were enriched within the expanded GC subsets in αv-villin mice, demonstrating concordant effects on GCs across these experimental systems.

The observed expansion of GC populations following αvβ6 disruption may appear discordant with the association of UC with GC depletion (*76, 77*). However, recent single-cell and spatial transcriptomic analyses indicate that UC involves dynamic remodeling of GC states rather than uniform lineage loss (*6, 41*). Consistent with this framework, the expanded GC subsets in αv-villin mice were enriched for antimicrobial and stress-associated genes and expressed *Gsdmc* family members, which are induced by type 2 cytokines (*78, 79*). Similar GC subsets are absent in germ-free mice and develop following microbial colonization and local immune cell activation (*57*), highlighting the influence of cytokine and microbial cues on goblet cell composition. Together, these findings support a model in which αvβ6-mediated TGFβ activation constrains cytokine- and microbiota-driven remodeling of goblet cell states.

Loss of epithelial αv also disrupted development of tissue-resident immune cells, with reduced numbers of some colonic IELs and near-complete loss of CD103 expression across all IEL subsets and intraepithelial DCs. CD103 expression is induced by TGFβ, strongly supporting a role for epithelial αv in locally activating TGFβ to sustain colon-resident immune populations. Reduced CD103^+^ mucosal T cells have been reported in active UC (*80, 81*), suggesting that impaired epithelial TGFβ activation phenocopies aspects of immune disruption seen in disease, and the similar IEL changes in the αvβ6-immunization model demonstrate that antibody-mediated blockade is sufficient to recapitulate these defects.

The concordance between autoantibody and gene knockout effects in mice supports our model that anti-αvβ6 antibodies contribute to disease through inhibition of integrin function rather than through immune targeting of epithelial cells (*82*). However, despite epithelial and immune alterations, neither epithelial αv deletion nor induction of αvβ6 autoantibodies triggered spontaneous inflammation. Both models did show increased inflammation and poor recovery in the DSS-colitis model. These results are consistent with prior studies of epithelial-intrinsic TGFβ disruption (*27–29*) and parallel the observation that individuals with αvβ6 autoantibodies may remain asymptomatic for years before UC diagnosis (*11*). We propose that αvβ6 dysfunction establishes a preclinical state of barrier vulnerability, characterized by expansion of potentially inflammatory GCs and tissue immune cells, which promote excessive inflammation.

These findings should be interpreted in light of several limitations. Although circulating anti-αvβ6 titers distinguished UC from healthy subjects, they showed weak associations with clinical activity and persisted after colectomy. Functional inhibition did not correlate directly with antibody titer, suggesting that qualitative features of the antibody repertoire, including epitope specificity and the fraction of function-blocking antibodies, may determine pathogenic potential. Additionally, DSS colitis represents an injury-driven model that does not fully recapitulate chronic human disease. Longitudinal studies in seropositive individuals, together with mechanistic analyses of the αvβ6 immunization model, will be necessary to define how anti-αvβ6 antibodies influence disease susceptibility and progression.

More broadly, this work reframes αvβ6 autoantibodies as functional modulators of epithelial signaling rather than mediators of direct epithelial immune targeting. By preventing localized activation of TGFβ at epithelial surfaces, these antibodies function as a de facto anti-cytokine response, attenuating a tissue-protective signaling pathway before overt inflammation develops. By linking preclinical seropositivity to impaired TGFβ activation and altered mucosal homeostasis, these findings provide mechanistic insight into how autoantibodies can shape disease susceptibility prior to overt inflammation. Therapeutic strategies aimed at restoring epithelial TGFβ signaling, preserving tissue-resident immune populations, or selectively neutralizing pathogenic anti-αvβ6 antibodies may therefore modify disease trajectory in at-risk individuals and potentially reduce severity once UC is established.

## Materials and Methods

### Study design

This study investigated whether UC-associated anti-αvβ6 autoantibodies inhibit epithelial TGFβ activation and signaling. Functional effects of αvβ6 blockade were examined in human colonic epithelial cells and *in vivo* using epithelial-specific αv deficient mice and induced anti-αvβ6 antibody models. DSS-colitis models were used to assess the impact of epithelial αvβ6 disruption on disease susceptibility. Human serum/plasma samples from UC and healthy subjects were analyzed for anti-integrin antibodies. Bulk RNA-seq datasets and human colon biopsy specimens were examined to assess gene expression and autoantibody production in inflamed versus noninflamed tissues. Sample sizes for each experiment are indicated in the corresponding figure legends.

### Human subjects

Participants were enrolled in either the Gastrointestinal Diseases or the Healthy Control Registry and Repository at the Benaroya Research Institute. Clinical protocols for each repository were approved by the BRI Institutional Review Board under protocol numbers IRB #10090 and IRB#3041700 respectively. All participants provided written informed consent prior to sample collection. Samples from healthy participants were age and sex matched to those with UC.

### Recombinant proteins

Recombinant human αvβ6 protein and anti-αvβ6 monoclonal antibodies 3G9 (Sequence from US patent US8992924B2) and F4 (*83*) were produced by WuXi Biologics. 3G9 was produced in a human IgG1 backbone while F4 was produced both in a human IgG1 and IgA backbone.

### ELISA

Immulon 2HB plates were coated with recombinant human or mouse αvβ6 (WuXi Biologics; R&D Systems) or human αvβ3 (R&D Systems) diluted in coating buffer. Diluted human or mouse serum/plasma or undiluted biopsy supernatants were added to coated plates, followed by incubated with HRP-conjugated secondary antibodies. Plates were developed with TMB substrate and absorbance was measured at 450 nm with background correction at 650 nm.

Seropositivity thresholds were defined using ROC analysis with Youden’s index. Quantitative antibody levels were determined using monoclonal standard curves. Total IgG and IgA were measured using commercial kits according to manufacturer instructions.

### ELISPOT

IgG and IgA ELISPOT assays were performed on PBMCs or lamina propria cells using recombinant αvβ6-coated PVDF plates. PBMCs were pre-activated prior to plating. Spots were developed according to manufacturer protocols and quantified using CellProfiler.

### Cell adhesion assays

Cell adhesion assays were performed using plates coated with fibronectin or latent TGFβ. HT-29 cells were pre-incubated with serum, purified IgG, or blocking antibodies and allowed to adhere. Adherent cells were quantified using a BioTek Cytation 3 imaging system. Inhibition was calculated relative to healthy subject controls.

Percent inhibition was calculated as:

(1 − [mean adherence of UC sample / mean adherence of healthy subjects]) × 100.

Individual UC samples were compared to the HS mean using one-sided Welch’s t tests. Samples were classified as inhibitory if percent inhibition was ≥30% and P ≤ 0.05.

### IgG purification and TGF**β** activation assays

IgG was purified from serum and tested for inhibition of αvβ6-mediated TGFβ activation using co-culture and monoculture reporter assays. TGFβ activation was assessed using HEK-Blue reporter cells (InvivoGen) plated on LAP-coated wells. Reporter activity was measured by SEAP production.

αvβ6-expressing reporter cells were generated by transfecting HEK-Blue cells with β6 pcDNA1neo (Addgene plasmid #13580) using Lipofectamine 3000. Stable transfectants were selected with G418 and sorted for high surface αvβ6 expression by flow cytometry.

### Culture of human IECs for RNA-Seq

T84 cells were differentiated on transwell inserts and treated with anti-αvβ6 (3G9), anti–TGF-β, active TGFβ, or media control prior to RNA isolation. Primary human intestinal epithelial cells (IECs) were derived from colonoscopic biopsies and cultured in a stem/progenitor-supporting system as previously described (55), with minor modifications. Primary IEC cultures were treated with 3G9, TGFβ, or media control prior to RNA stabilization and extraction. Additional culture conditions and processing steps are provided in Supplementary Methods.

### RNA-Sequencing

RNA libraries were prepared from purified RNA and sequenced on an Illumina platform. Reads were aligned to the GRCh38 reference genome and gene counts generated. Differential expression was performed using the limma-voom framework with Benjamini–Hochberg correction. Quality control metrics are described in Supplementary Methods.

### Differential gene expression analysis of human IECs

Differential gene expression was analyzed using limma-voom with empirical Bayes moderation. Primary IEC analyses incorporated donor pairing as a blocking factor. Principal component analysis and pathway enrichment were performed in R. Gene ontology enrichment was assessed using clusterProfiler with FDR correction.

### Colon epithelial marker analysis

Log2-transformed normalized RNA-seq expression values were used to quantify epithelial differentiation programs using curated marker gene sets representing major intestinal epithelial subsets. For the T84 dataset, gene-wise Z-scores were calculated across samples, and subset signature scores were computed as the mean Z-scored expression of marker genes per sample. For the primary IEC dataset, which contained heterogeneous epithelial populations, gene set enrichment was quantified using Gene Set Variation Analysis (GSVA) on filtered log2-transformed expression values. Signature scores were visualized as boxplots and heatmaps of group-mean values. Marker gene expression patterns were visualized using Z-score–normalized heatmaps. All analyses were performed in R.

### PC loading and pathway analysis

Principal component analysis (PCA) was performed on log-transformed, normalized gene expression matrices in R. Genes with the highest positive or negative loadings for principal components of interest were used for heatmap visualization and pathway enrichment analysis. Gene Ontology (GO) Biological Process enrichment was performed using clusterProfiler with Benjamini–Hochberg correction. Full analytical parameters are provided in Supplementary Methods.

### Mouse models

Intestinal epithelium-specific αv-knockout mice (αv-villin) were generated by crossing *Itgav*^flox/flox^ mice (*84*) on a C57Bl/6 background and Vil1-cre mice (B6.Cg-Tg(Vil1-cre)1000Gum/J; Jackson Laboratory). Littermate *Itgav*^flox/flox^ mice with no Cre transgene were used as controls. αv-villin mice of 8-16 weeks were used for experiments. Immunizations were performed on C57BL/6J mice (Jackson laboratory) at 6-8 weeks of age. All mice were housed under specific pathogen-free conditions at Benaroya Research Institute. All animal experiments were performed under appropriate licenses and institutional review within local and national guidelines for animal care.

For induction of αvβ6 autoantibodies, C57BL/6J mice (Jackson Laboratory) were immunized subcutaneously with recombinant human αvβ6 protein (WuXi Biologics) emulsified in Complete Freund’s Adjuvant (CFA; InvivoGen), followed by booster immunization in Incomplete Freund’s Adjuvant (IFA; InvivoGen) two weeks later. Serum anti-αvβ6 responses were confirmed by ELISA. IEL analysis was performed three months after boost, and DSS colitis experiments were initiated one month post-boost.

### DSS colitis

Mouse colitis was induced by administration of 1.75 – 2% w/v of dextran sodium sulfate (DSS; MP Biomedicals, 160110) *ad libitum* dissolved in drinking water for 7 days, followed by 7 days of regular water. General health and body weight were monitored every other day to daily. Mice that lost more than 20% of starting weight were euthanized. Severity of colitis was assessed using a disease activity index (DAI) which incorporates weight loss, stool consistency, and GI bleeding (*85*). On day 14, colons were harvested and lengths recorded. Colon swiss rolls were prepared as described below, stained with hematoxylin and eosin (H&E), and samples were scanned on an ImageXpress HTai confocal microscope using 10X objective.

### Generation of colon swiss rolls for spatial transcriptomics and RNAscope

Murine colons were excised, opened longitudinally, cleaned of luminal contents, and rolled from distal to proximal to generate swiss roll preparations. Tissues were fixed, paraffin-embedded, and sectioned for spatial transcriptomics or RNAscope analysis. Spatial transcriptomic libraries were generated using the 10x Genomics Visium HD platform according to manufacturer protocols.

### *in situ* hybridization

RNAscope was performed using the RNAscope Multiplex Fluorescent Assay (Advanced Cell Diagnostics) according to manufacturer instructions. FFPE colon swiss rolls were processed according to the manufacturer’s protocol, sectioned at 10-micron thickness, and hybridized with four target-specific probes: *Duoxa2, Gsdmc4, Aqp8*, and *Retnlb*. Signals were developed using TSA fluorophores, nuclei were counterstained with DAPI, and slides were imaged using identical acquisition settings across samples on an Akoya Biosciences fluorescence imaging system.

### Spatial transcriptomics analysis

Spatial transcriptomics was performed on FFPE colon swiss rolls using the 10x Genomics Visium HD platform. Libraries were prepared and sequenced according to manufacturer protocols. Data were processed using Space Ranger and analyzed in Python using Scanpy-based workflows. Cell segmentation, clustering, and trajectory analyses were performed as described in Supplementary Methods.

### Mouse intestinal preparation for flow cytometry

Murine colon and cecum were excised, opened longitudinally, and washed to remove luminal contents. Intraepithelial lymphocytes (IELs) were isolated using sequential epithelial stripping and density gradient separation. Remaining tissue was enzymatically digested to obtain lamina propria lymphocytes (LPLs). Single-cell suspensions were filtered, washed, and processed for flow cytometric analysis.

### Flow cytometry

Intraepithelial and lamina propria lymphocytes were isolated from murine colon and analyzed by flow cytometry using standard enzymatic digestion and density gradient separation. Cells were stained with viability dye and surface markers, fixed, and analyzed on a BD Fortessa or Symphony cytometer. Gating strategies are shown in Supplementary Fig. 9.

### Statistics

Statistical analyses were performed using GraphPad Prism (v10-11) unless otherwise indicated. RNA-Seq analyses were conducted in R. All tests were two-tailed unless otherwise specified, and P < 0.05 was considered statistically significant. Significance is denoted as ns (P > 0.05), *P < 0.05, **P < 0.01, ***P < 0.001, ****P < 0.0001. Two-group comparisons were performed using unpaired two-tailed t tests with Welch’s correction or Mann-Whitney U tests for nonparametric data. For the adhesion assays, one-sided Welch’s t tests were performed comparing individual UC samples to the healthy subject mean. Correlations were assessed using Spearman rank correlation. ROC curves were generated in Prism, and optimal thresholds were defined by maximizing Youden’s index. Multiple comparisons were adjusted using the two-stage step-up false discovery rate (FDR) method where indicated. Survival was analyzed using Kaplan–Meier curves and the log-rank (Mantel-Cox) test. Longitudinal data were analyzed using mixed-effects models (REML). RNA-Seq differential expression was performed using the limma-voom framework with Benjamini–Hochberg FDR correction. Paired primary IEC analyses incorporated duplicate correlation to model donor effects.

## Acknowledgments

We would like to thank the participants in Benaroya Research Institute (BRI) Registry and Repository (RRID: SCR_026967) who donated blood and/or tissue for this study. We acknowledge the BRI Center for Interventional Immunology and BRI Clinical Core Laboratory for blood and biopsy collection and processing, the BRI Animal Resources Core, the BRI Cell and Tissue Analysis Core (RRID: SCR_026327) for flow cytometry, histology, and microscopy, and the BRI Genomics Core (RRID: SCR_026658) for RNA-seq and 10X visium. We also thank the M.J. Murdock Charitable Trust for generously providing equipment funding for the BRI Cores. We would also like to thank the Histology and Imaging Core at Fred Hutchinson Cancer Center for assistance with RNAscope (supported by NIH P30 CA015704 of the Fred Hutch/University of Washington/Seattle Children’s Cancer Consortium). Figures 2D, 4A, and Supp Fig 2D were created using Biorender.com. Artificial intelligence (ChatGPT, OpenAI; GPT-5.2) was used to assist with grammar refinement and generation of preliminary RNA-seq analysis scripts. All scientific content, analyses, and interpretations were independently verified by the authors.

## Funding

Research reported in this publication was supported by funding from the National Institutes of Health (70% of funding) and from non-governmental sources (30%), listed below:

National Institutes of Health/ NIDDK grant R01DK136071 (to ALH).

National Institutes of Health/ NIAID grant R21AI171921 (to ALH and OJH)

Kenneth Rainin Foundation research grant KRA000163 (ALH and JDL)

McCaw Family Foundation grant (CS)

Research infrastructure at BRI was supported in part by grants from the Murdock Trust.

The content is solely the responsibility of the authors and does not necessarily represent the official views of the National Institutes of Health.

## Author contributions

Conceptualization: KJF, ALH

Data curation: KJF, AEY, SS, AM, MES, JDL

Formal analysis: KJF, LWT, THE

Funding acquisition: CS, JDL, ALH

Methodology: KJF, SS, AM, MES, CS, JDL

Investigation: KJF, AEY, JFM, KEM, DMS, DGK, JDL, ALH

Visualization: KJF, KEM, LWT, THE, CS, DGK, ALH

Supervision: KJF, ALH

Writing – original draft: KJF, ALH

Writing – review & editing: KJF, JFM, KEM, LWT, THE, CS, OJH, CS, JDL, ALH

## Competing interests

Authors declare that they have no competing interests.

## Data and materials availability

In progress.

## Supplementary Materials

Fig. S1: Serum antibody profiling and clinical associations in ulcerative colitis Fig. S2: Validation of αvβ6-dependent adhesion and activation of latent TGFβ

Fig. S3: Differential expression and pathway enrichment analyses in TGFβ-perturbed IECs Fig. S4: Representative histology and colon lengths of control and αv-villin mice

Fig. S5: Spatial transcriptomics of re-clustered GCs and enterocytes Fig. S5: Spatial transcriptomics of re-clustered GCs and enterocytes

Fig. S6: Baseline immune cell composition in unchallenged control and αv-villin mice Fig. S7: Representative H&E of tissue from immunized mice

Fig. S8: ‘Digital unrolling’ of colon swiss rolls

Fig. S9: Gating strategy for intraepithelial immune cells

Table S1: Curated human intestinal epithelial marker gene sets Table S2: Flow cytometry antibodies used for mouse phenotyping Data file S1: Differential gene expression in treated T84

Data file S2 Differential gene expression in treated primary IEC

## Notes

### Competing Interest Statement

The authors have declared no competing interest.

### Summary of Updates

The author order has been updated to list Kayla Fasano as first author and Adam Lacy-Hulbert as senior author. No changes were made to the manuscript files

