## Supplemental Figures for "UC-associated autoantibodies to αvβ6 inhibit mucosal TGFβ activation and predispose to intestinal inflammation"

This file includes:

Figures S1 to S9

Tables S1 to S2

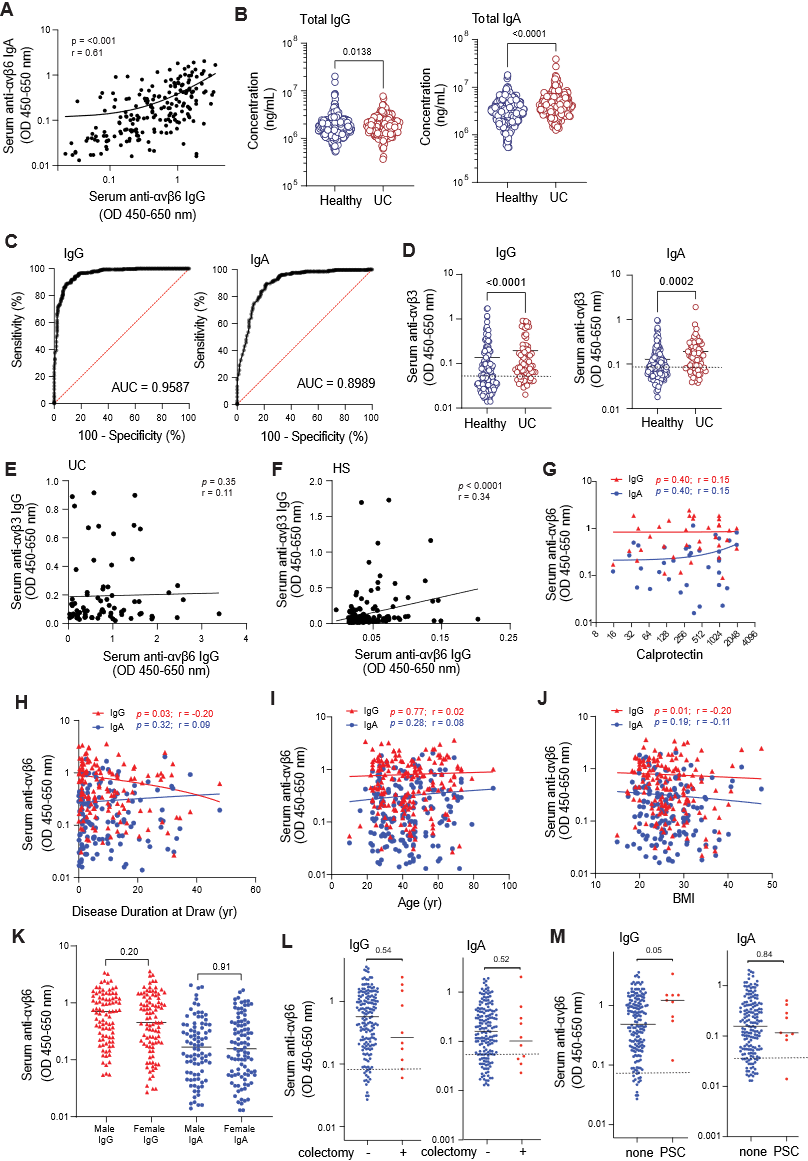

**Fig. S1: Serum antibody profiling and clinical associations in ulcerative colitis. (A)** Correlation between serum anti-αvβ6 IgG and IgA OD in UC. **(B)** Total serum IgG (left) and IgA (right) quantified by ELISA in HS (n=338) and UC (n = 194). **(C)** Receiver operating characteristic (ROC) curves differentiating UC from controls based on serum anti-αvβ6 IgG (left) and IgA (right) ELISA OD, with areas under the curve (AUC) shown. **(D)** Serum anti-αvβ3 IgG (left) and IgA (right) were quantified by ELISA in HS (n = 157) and UC (n = 76). **(E,F)** Correlation of serum anti-αvβ6 and anti-αvβ3 IgG in UC **(E)** or HS **(F)**. **(G-J)** Correlation of IgA (blue) and IgG (red) OD values in UC with contemporaneous fecal calprotectin (n=35) **(G)**, duration of disease at sample collection **(H)**, age **(I)**, and body mass index (BMI, n=154) **(J)**. **(K)** Anti-αvβ6 IgG (red) or IgA (blue) by sex in UC. **(L,M)** anti-αvβ6 OD for IgG (left) and IgA (right) for UC who had (n=10; red) or had not (n=173; blue) previously undergone curative colectomy **(L)**, or UC with (n=9; red) or without (n=174; blue) primary sclerosing cholangitis (PSC) **(M)** Dashed lines indicate OD thresholds for autoantibody positivity, determined by receiver operating characteristic (ROC) curve analysis using the highest Youden’s index as shown in **Fig 1**. Data are shown as individual data points with medians and two-group comparisons were performed using two-tailed Mann-Whitney tests **(B, D, K–M)**. Correlations were assessed using two-tailed Spearman rank correlation, and best-fit lines were included for visualization purposes only **(A, E-J)**.

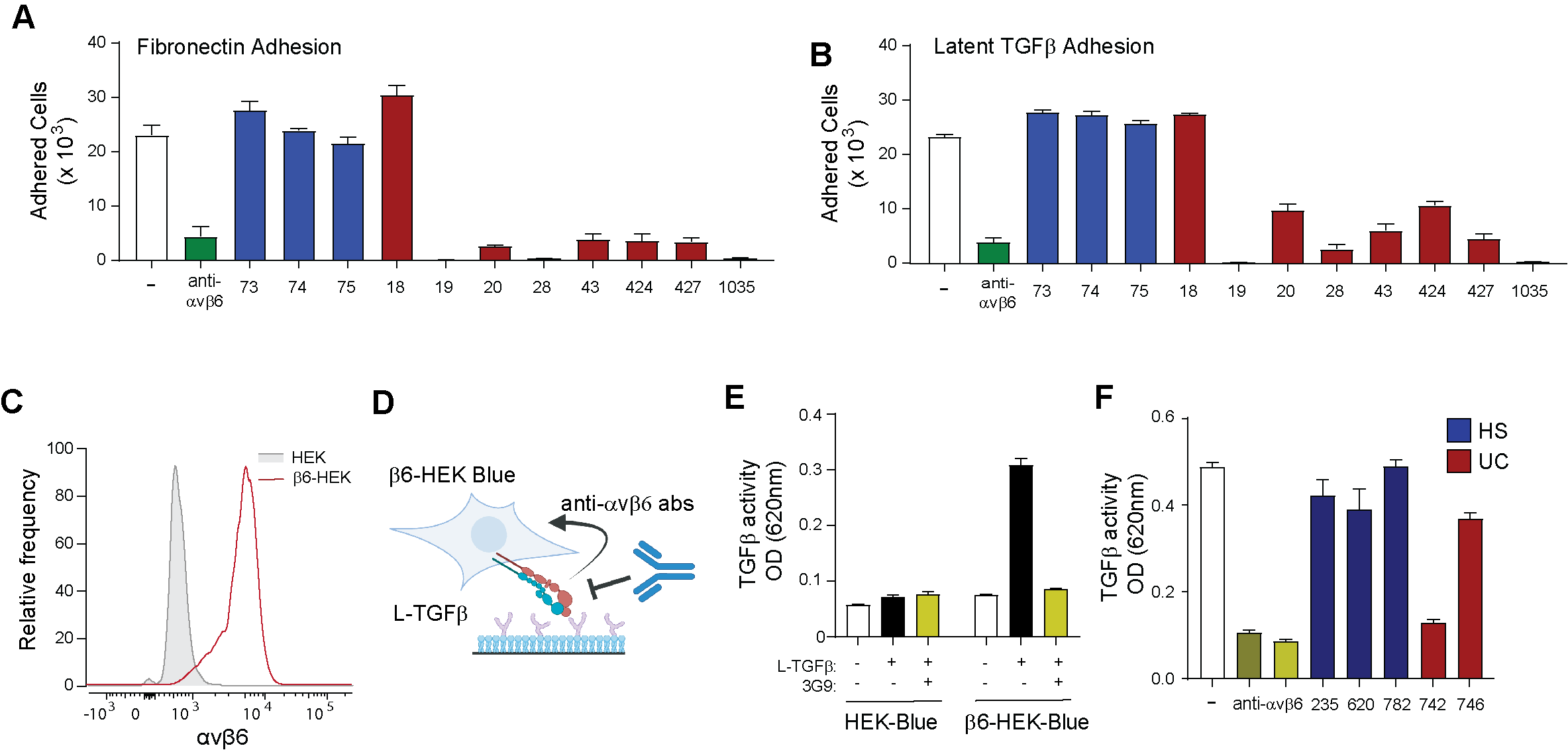

**Fig. S2: Validation of αvβ6-dependent adhesion and activation of latent TGFβ**. **(A-B)** Adhesion of HT-29 cells to fibronectin **(A)** or L-TGFβ **(B)** in the presence of 3G9 (green) or sera from HS (blue) and UC (red). **(C)** Representative flow cytometry histograms showing surface expression of αvβ6 on parental HEK-Blue cells (gray) and β6-transfected HEK-Blue cells (β6-HEK-Blue; red). **(D)** Schematic of the β6-HEK-Blue L-TGFβ activation assay. **(E)** Activation of L-TGFβ by parental and β6-HEK-Blue cells ± 3G9. **(F)** HEK-Blue reporter activity following incubation with L-TGFβ in the presence of purified IgG from HS and UC. Data are shown as mean ± SEM.

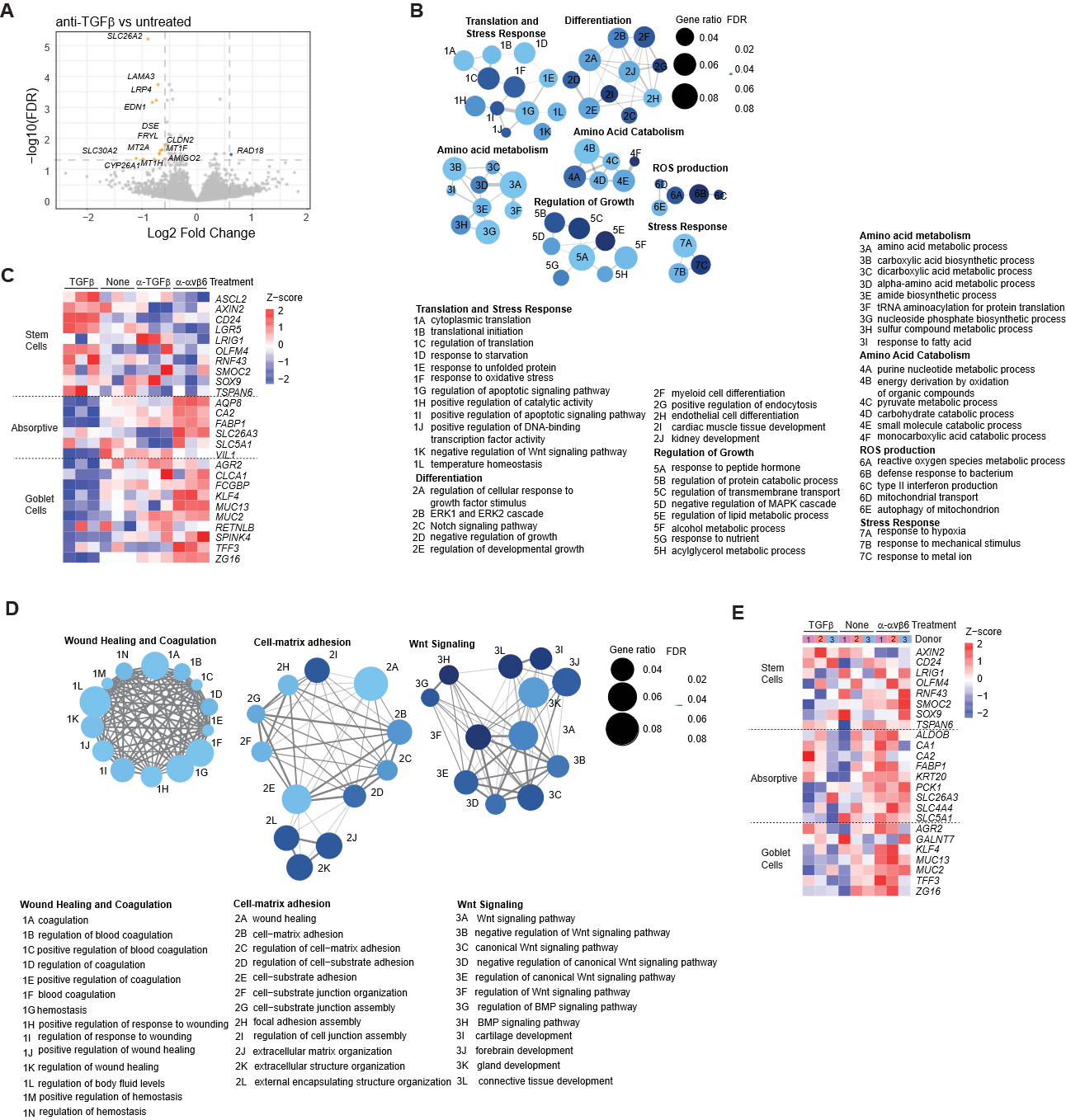
**Fig. S3: Differential expression and pathway enrichment analyses in TGFβ-perturbed IECs. (A)** Volcano plot of DEGs in T84 cells treated with anti-TGFβ versus untreated. **(B)** Network representation of Gene Ontology (GO) Biological Process terms enriched among positive PC1 loadings (TGFβ-associated response) in T84 cells. **(C)** Heatmap of individual marker genes comprising the epithelial subset signature scores in T84 cells from **Fig 3E**. **(D)** Network representation of GO Biological Process terms enriched among genes with positive PC2 loadings (TGFβ-associated response) in primary IECs. **(E)** Heatmap of individual marker genes used for GSVA-based epithelial program enrichment in primary IECs from **Fig 3I**. Dotted lines in volcano plot indicate fold-change and FDR thresholds.

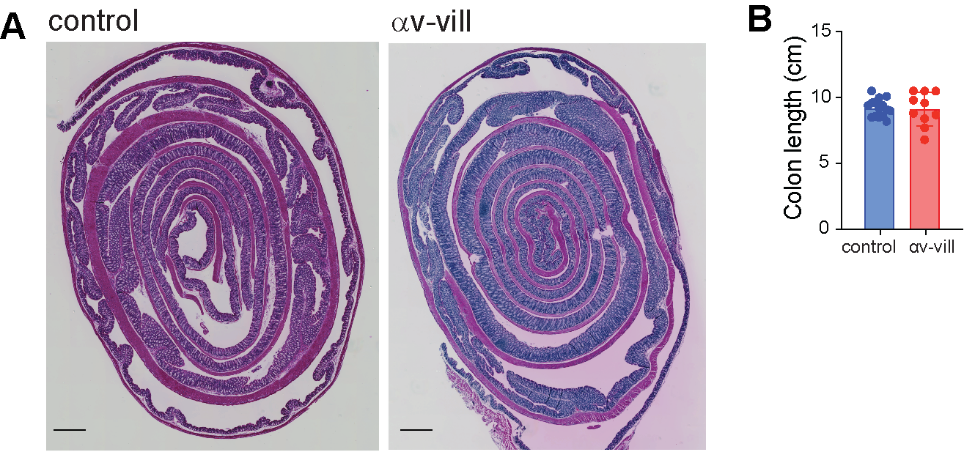

**Fig. S4: Representative histology and colon lengths of control and αv-villin mice. (A)** H&E stained swiss roll sections of whole colon from representative control and αv-villin mice (captured at 4X, scale bar 1mm). **(B)** Colon length in naïve control and αv-villin mice at baseline.

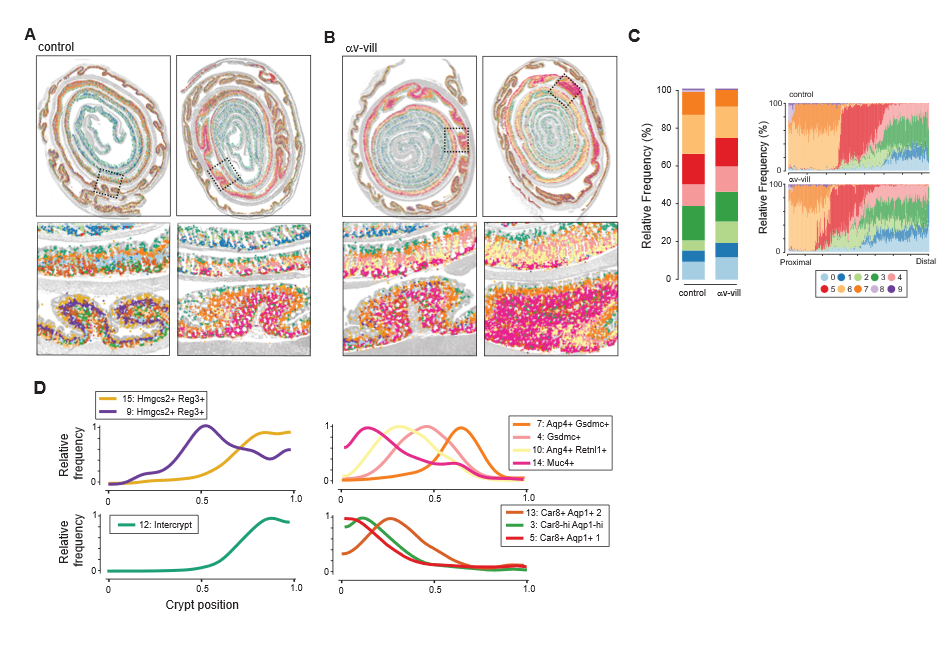
**Fig. S5: Spatial transcriptomics of re-clustered GCs and enterocytes. (A-B)** Spatial projection of GC clusters over H&E image (grayscale) (4X magnification, scale bar 1mm). Each image is from an individual mouse. Regions within dashed lines are shown in higher magnification below each image. **(C)** Relative proportions of enterocyte clusters in control and αv-villin mice, as colored in **Fig 5E**. Left panel shows relative frequency across the whole colon, right panel shows frequencies for sequential regions along the length of the colon. **(D)** Distribution of indicated GC clusters described in **Fig 5F** along the crypt-lumen axis (combined data from 2 control and 2 αv-villin mice).

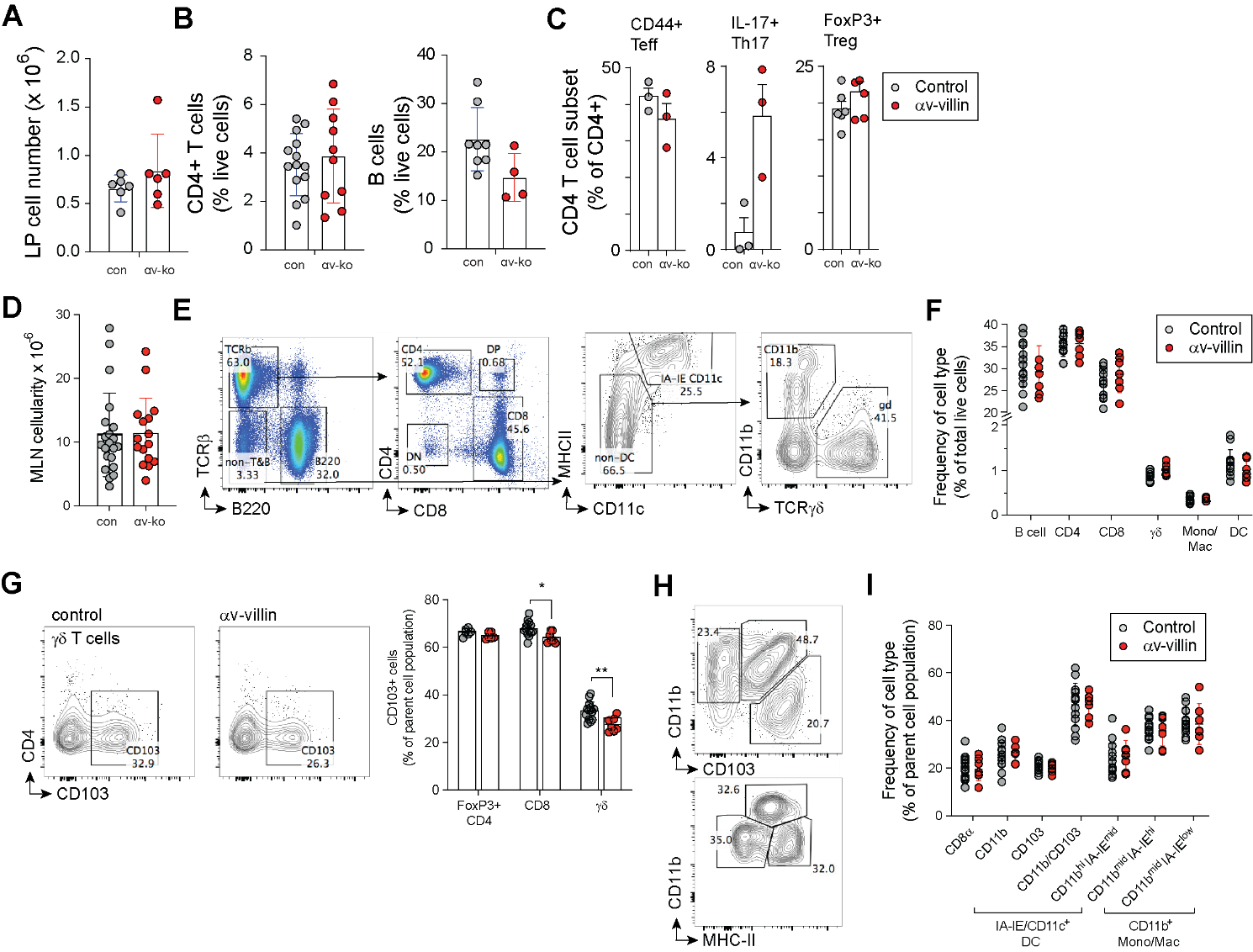

**Fig. S6: Baseline immune cell composition in unchallenged control and αv-villin mice. (A-C)** Quantification of total lamina propria (LP) cells **(A)** and frequencies **(B)** of LP CD4⁺ T cells (left) and B cells (right). **(C)** Frequencies of T cell subsets among CD4⁺ T cells. **(D-I)** Quantification of mesenteric lymph node (MLN) cells. (F) Total MLN cellularity, with **(G)** representative gating for lymphoid and myeloid populations. **(H)** Frequencies of immune cell subsets. **(I)** Representative plots and quantification of CD103⁺ cells among T cells subsets. **(J)** Gating strategy for intestinal myeloid subsets and **(K)** frequencies of indicated populations among parent gates. Data are shown as mean ± SD. Statistical significance was determined using unpaired t tests with Welch’s correction. * = P < 0.05; ** = P < 0.01.

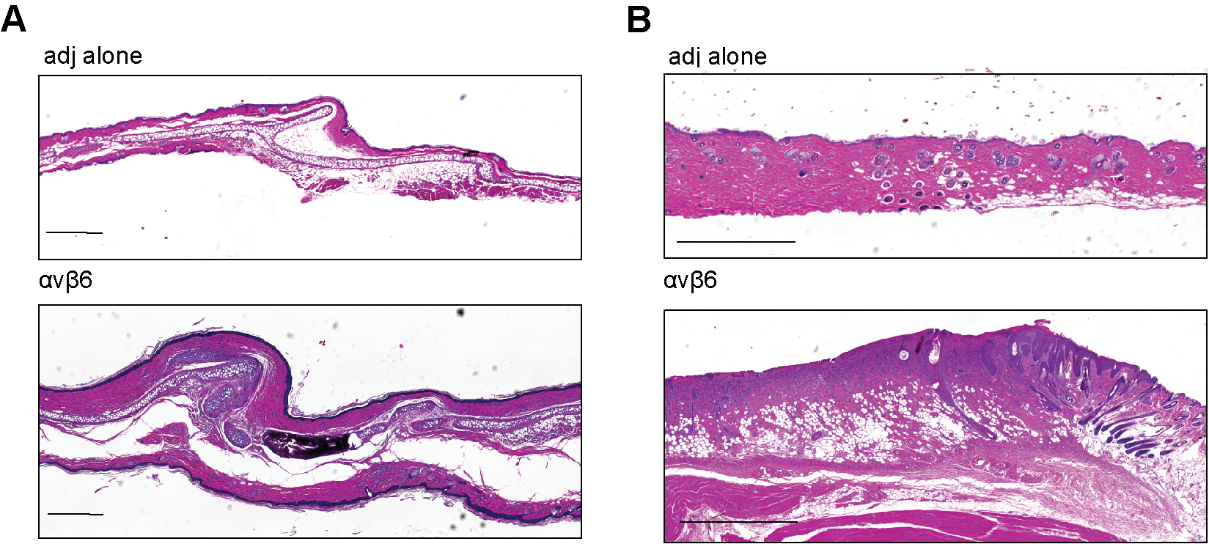

**Fig. S7: Representative H&E of tissue from immunized mice. (A-B)** Representative H&E of ear **(A)** or skin at site of immunization **(B)** from αvβ6-immunized and adjuvant-only mice collected 89 days post-boost. Scale bars 500μM.

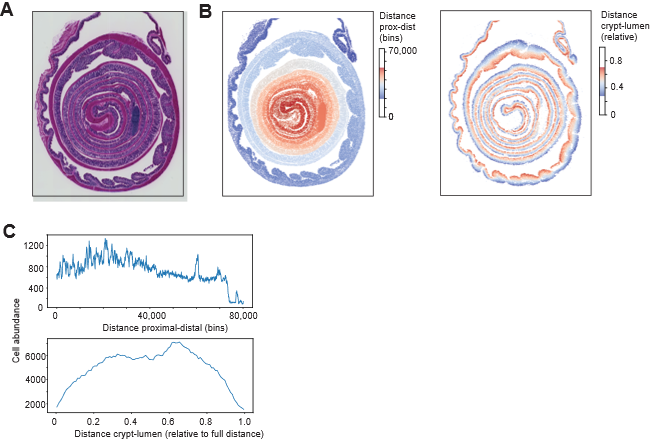

**Fig. S8: ‘Digital unrolling’ of colon swiss rolls. (A-C) (A)** H&E stain of representative section (4X magnification), and **(B)** cells colored based on calculated position along the proximal to distal axis (left) and enterocytes and GCs colored based on position in the crypt to lumen axis (right). **(C)** Histograms showing distribution of cells identified along the proximal – distal axis (top) or crypt–lumen axis (bottom).

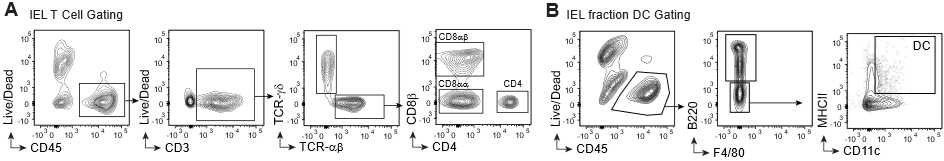

**Fig. S9: Gating strategy for intraepithelial immune cells. (A-B)** Representative gating strategy for identification of intraepithelial lymphocyte (IEL) T cell subsets **(A)** or IEL-associated DCs **(B)**.

| **Subset** | **Gene** |
| --- | --- |
| Stem | LGR5 |
| Stem | ASCL2 |
| Stem | OLFM4 |
| Stem | AXIN2 |
| Stem | RNF43 |
| Stem | SOX9 |
| Stem | SMOC2 |
| Stem | CD24 |
| Stem | TSPAN6 |
| Stem | LRIG1 |
| Absorptive | CA2 |
| Absorptive | CA1 |
| Absorptive | SLC26A3 |
| Absorptive | AQP8 |
| Absorptive | ALDOB |
| Absorptive | FABP1 |
| Absorptive | SLC9A3 |
| Absorptive | APOA1 |
| Absorptive | VIL1 |
| Absorptive | SLC5A1 |
| Goblet | MUC2 |
| Goblet | TFF3 |
| Goblet | FCGBP |
| Goblet | SPINK4 |
| Goblet | MUC13 |
| Goblet | AGR2 |
| Goblet | ZG16 |
| Goblet | RETNLB |
| Goblet | CLCA1 |
| Goblet | KLF4 |

**Table S1: Curated human intestinal epithelial marker gene sets.** Marker genes representing major human colonic epithelial subsets were defined based on published intestinal epithelial atlases and used for signature and GSVA analyses.

| T Cell Panel (Immunized) - Fortessa | | | |
| --- | --- | --- | --- |
| Marker | Fluorophore | Supplier | Catalog Number |
| CD45 | BUV395 | BD | 564279 |
| Fixable Viability Stain | 440UV | BD | 566332 |
| CD4 | BUV737 | BD | 612844 |
| CD11c | e450 | ThermoFisher | 48-0114-80 |
| CD103 | BV510 | BioLegend | 121423 |
| CD8ß | BV605 | BD | 740387 |
| Thy1.2 | BV786 | BioLegend | 105331 |
| KLRG1 | FITC | BioLegend | 138409 |
| TCRß | PerCP-Cy5.5 | ThermoFisher | 45-5961-82 |
| CD3 | PE | BD | 553064 |
| CD44 | PE-Cy7 | BioLegend | 103029 |
| CD69 | APC | BioLegend | 104513 |
| CD8α | AF700 | ThermoFisher | 56-0081-80 |
| T Cell Panel (αV-Villin) - Symphony | | | |
| Marker | Fluorophore | Supplier | Catalog Number |
| CD45 | BUV395 | BD | 564279 |
| CD4 | BUV805 | BD | 569193 |
| TCRγδ | BV421 | BD | 562892 |
| CD103 | BV510 | BD | 563087 |
| CD8ß | BV605 | BD | 740387 |
| CD69 | BV785 | BioLegend | 104543 |
| FoxP3 | FITC | ThermoFisher | 11-5773-82 |
| TCRß | PerCP-Cy5.5 | ThermoFisher | 45-5961-82 |
| CD3 | PE | BD | 553064 |
| CD44 | PE-Cy7 | BioLegend | 103029 |
| CD8α | AF700 | ThermoFisher | 56-0081-80 |
| Fixable Viability Stain | Zombie NIR | BioLegend | 423105 |
| DC Panel (αV-Villin) - Symphony | | | |
| Marker | Fluorophore | Supplier | Catalog Number |
| CD45 | BUV395 | BD | 564279 |
| B220 | BUV737 | BD | 612838 |
| Fixable Viability Stain | Violet | ThermoFisher | L34964 |
| CD103 | BV510 | BD | 563087 |
| CD24 | BV650 | BD | 563545 |
| CD4 | BV786 | BD | 563331 |
| MHCII | FITC | BD | 553623 |
| CD11b | PerCP-Cy5.5 | BD | 550993 |
| CD8α | PE | BD | 553033 |
| F4/80 | PE-Cy7 | BioLegend | 123114 |
| CD11c | APC | BD | 550261 |

**Table S2: Flow cytometry antibodies used for mouse phenotyping**
